# SunCatcher: Clonal Barcoding with qPCR-Based Detection Enables Functional Analysis of Live Cells and Generation of Custom Combinations of Cells for Research and Discovery

**DOI:** 10.1101/2021.10.13.464251

**Authors:** Qiuchen Guo, Milos Spasic, Adam Maynard, Gregory J. Goreczny, Jessica F. Olive, Amanuel Bizuayehu, Sandra S. McAllister

**Affiliations:** Division of Hematology, Department of Medicine, Brigham and Women’s Hospital, Boston, MA 02115, USA; Department of Medicine, Harvard Medical School, Boston, MA 02115, USA; Broad Institute of Harvard and MIT, Cambridge, Massachusetts, 02142, USA; Harvard Stem Cell Institute, Cambridge, Massachusetts, 02138, USA

## Abstract

Over recent decades, cell lineage tracing, clonal analyses, molecular barcoding, and single cell “omic” analysis methods have proven to be valuable tools for research and discovery. Here, we report a clonal molecular barcoding method, which we term SunCatcher, that enables longitudinal tracking and retrieval of live barcoded cells for further analysis. Briefly, single cell-derived clonal populations are generated from any complex cell population and each is infected with a unique, heritable molecular barcode. One can combine the barcoded clones to recreate the original parental cell population or generate custom pools of select clones, while also retaining stocks of each individual barcoded clone. We developed two different barcode deconvolution methods: a Next-Generation Sequencing method and a highly sensitive, accurate, rapid, and inexpensive quantitative PCR-based method for identifying and quantifying barcoded cells *in vitro* and *in vivo*. Because stocks of each individual clone are retained, one can analyze not only the positively selected clones but also the negatively selected clones result from any given experiment. We used SunCatcher to barcode individual clones from mouse and human breast cancer cell lines. Heterogeneous pools of barcoded cells reliably reproduced the original proliferation rates, tumor-forming capacity, and disease progression as the original parental cell lines. The SunCatcher PCR-based approach also proved highly effective for detecting and quantifying early spontaneous metastases from orthotopic sites that would otherwise have not been detected by conventional methods. We envision that SunCatcher can be applied to any cell-based studies and hope it proves a useful tool for the research community.

## Introduction

Cell fate mapping and lineage tracing techniques have been employed for several decades and have been useful for describing heterogeneity and clonal dynamics of complex tissues in both normal and pathological settings. Early insights into hematopoiesis were derived from analyzing karyotypes, assessing heterozygosity of particular genes, or analyzing viral integration patterns^1-3^. Although limited by their lack of sensitivity and their qualitative nature, those techniques enabled the first effective tracking of clonal populations.

Understanding cellular heterogeneity and clonal fitness has proven particularly important to cancer research. Clinically, intratumoral diversity inversely correlates with therapeutic response, metastatic ability, and patient survival^4-6^. Although differences in the mutational profile of tumor cell subpopulations may be the best-documented parameter of tumor heterogeneity, it is ultimately the functional heterogeneity, a result of both genetic and non-genetic sources of heterogeneity, that impacts the course of disease progression^7-10^. Large scale sequencing efforts and technologies such as single cell RNA sequencing and single nucleus sequencing have revealed the diverse mutational and transcriptional landscape of human cancers^11-15^. Nevertheless, we still have much to learn about the effects of functional heterogeneity on disease progression.

Accounting for and properly modeling tumor heterogeneity and cell plasticity in the experimental and pre-clinical settings is thus crucial for improving cancer treatment outcomes. For example, we previously reported that functional heterogeneity can impact data interpretation in studies that employ gene editing and designed a novel CRISPR/Cas9 protocol that incorporates a single cell cloning step prior to gene editing, in order to generate appropriately matched control and edited cells^16^. Moreover, clonal dynamics–driven by clone frequency, spatial proximity, heterotypic interactions, and/or cell plasticity–can profoundly affect disease progression^4, 10, 17, 18^.

The advent of molecular barcoding techniques, in which short, non-coding DNA sequences are stably integrated into the genomes of complex cell populations using viral infection, enables one to track thousands of clonal populations^19-21^. In cancer research, DNA barcoding techniques have been used to successfully trace clonal dynamics among heterogeneous tumor cell populations^22-29^. Those tracing techniques utilize barcode library infection methods in which 10^6^-10^8^ barcodes are introduced into a population of tumor cells. While such high-complexity barcode libraries enable longitudinal quantification of clonal diversity in various experimental settings, they do not enable characterization, functional analyses, or manipulation of the cellular populations of interest. More recently, a number of elegant and sophisticated methods have been developed to trace as well as study barcoded cells that comprise tumors and contribute to disease progression and therapeutic resistance^30-34^.

Here, we report a novel molecular barcoding approach, termed SunCatcher, that provides a rapid, inexpensive, and highly sensitive method to detect, identify, quantify, isolate, and study individual clones, or mixtures of clones, of interest. The SunCatcher approach is based on generating single cell-derived clonal populations from any complex cell population and infecting each with a unique, heritable molecular barcode. The barcoded clones can be studied individually, combined to recreate the original parental cell population, or combined to generate custom pools of select clones. As such, SunCatcher enables both longitudinal and retrospective analysis of the molecular, phenotypic, and functional basis for heterogeneity and clonal fitness during experimentation. Importantly, our approach even enables one to study clones that are negatively selected during any process so that their specific vulnerabilities can be uncovered.

We applied SunCatcher to barcode individual clones from a number of mouse and human breast cancer cell lines. Heterogeneous pools of barcoded cells reliably reproduced the original growth and tumor progression rates–in vitro and in vivo–as the original parental cell lines. The SunCatcher approach also proved highly effective for detecting and identifying early spontaneous skeletal and visceral metastases. We envision that SunCatcher can be applied to any cell-based studies and hope it proves useful for research and discovery.

## Results

### SunCatcher Clonal Barcoding Approach

We developed a clonal barcoding system, which we call SunCatcher, that enables longitudinal and retrospective analyses of live cells of interest. First, single cells are isolated from a heterogeneous parental population of cells, by either dilution passage or fluorescence-activated cell sorting (FACS) based on cell-surface markers of interest, into individual wells of 96-well plates (Fig. 1A). Each single cell-derived colony is then expanded, after which aliquots of cells from each clonal population are viably frozen to generate stocks of non-barcoded clones (NBCs).

**Figure 1.**
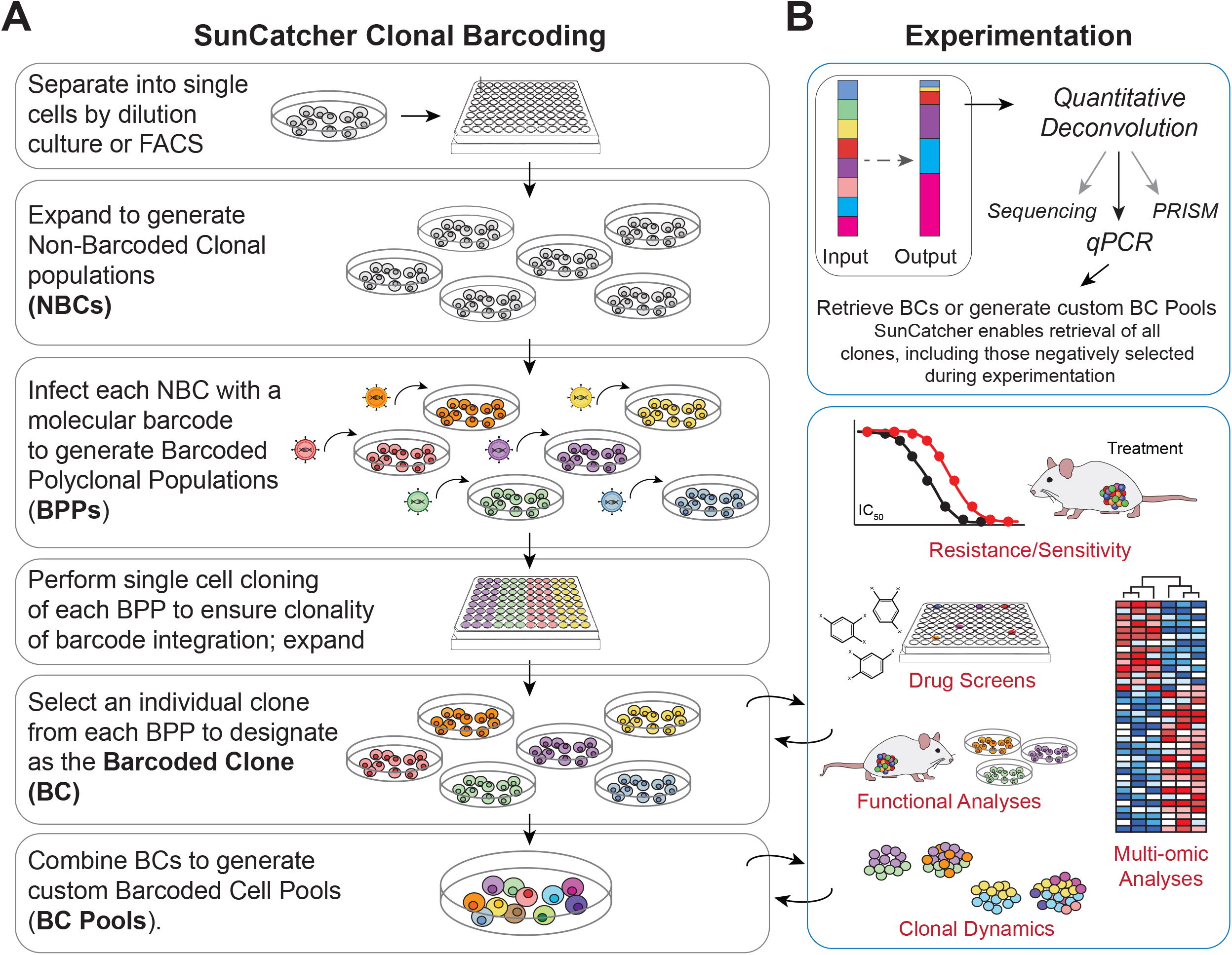
SunCatcher Clonal Barcoding and Functional Analyses. (**A**) Schematic of the SunCatcher Clonal Barcoding Method. SunCatcher utilizes two rounds of single cell cloning to ensure that each subclone has only 1 unique barcode and that each cell within that subclone contains the same barcode insertion site. Due to the single cell cloning approach, custom BC pools of any combination can be designed. (FACS, fluorescence-activated cell sorting; NBC, non-barcoded clone; BPP, barcoded polyclonal population (polyclonal for barcode insertion site); BC, barcoded clone; BC Pool, population containing multiple BCs). (**B**) BCs or BC Pools (input) can be entered into any experiment and detected at end point (output) using various deconvolution approaches. Deconvolution by qPCR provides a cost effective, rapid, and highly sensitive method for detecting and quantifying BCs. SunCatcher enables retrieval of all clones, including those negatively selected during experimentation, for further analyses and/or design and testing of custom BC Pools.

Each NBC population is then infected with a unique, heritable molecular barcode (Fig. 1A); in this case, we used the PRISM library of lentiviral vectors that each contains a unique, heritable 24-base-pair (bp) DNA barcode sequence designed to avoid sequence homology with the genome^28, 35^. To ensure clonality of the barcode insertion site, we perform a subsequent round of single cell cloning of each barcoded polyclonal population; those subclones are expanded and aliquots of each are viably frozen (Fig. 1A). Next, one population for each barcode is randomly selected to represent the barcoded clone (BC). At this stage, each BC has a single, unique 24-bp barcode and is clonal for the lentiviral insertion site. Each BC population is aliquoted into cryovials and viably frozen to generate BC stocks. The BCs can be analyzed individually or mixed in various combinations to create custom barcoded pools (BC Pools) to be used in any experiment or assay (Fig. 1A, B).

Quantitative analysis of barcode composition is performed by isolating genomic DNA followed by any number of deconvolution approaches, including PRISM analysis^28^, a next generation sequencing (NGS) assay that we developed, or our low-cost, rapid qPCR-based approach (Fig. 1B). The BC detection methods are described below, following the discussions of heterogeneity.

### Considering and Confirming Population Complexity when Generating Subclones

Estimating the number of clones required to represent the heterogeneity of a complex parental cell line is an important consideration in the SunCatcher approach. The SunCatcher clonal method can be applied to any cell population and so decisions about the number of clones required might be best made by considering the relevant phenotypes one wishes to capture.

Our own research is focused on breast cancer. While breast cancer subtypes are defined based on hormone receptor expression and amplification of the Her2/neu oncogene, our parental cell lines were each originally derived from tumors of a distinct subtype; the breast cancer cell lines we have barcoded to date include: the McNeuA Her2-positive^36^, 4T1 triple-negative breast cancer (TNBC)^37^, and Met1 TNBC^38^ murine mammary carcinoma cell lines, as well as the human HMLER-HR TNBC cell line^39^ (Table S1).

We made calculations based on reported analyses of breast tumor epithelium. Although complete understanding of the spectrum of heterogeneity and differentiation states of human breast epithelium and breast cancers is lacking, recently reported cell atlases provided helpful insights. For example, gene expression analysis had identified 6 molecular subtypes within the TNBC subtype^40^. A single cell RNAseq analysis of normal breast epithelium revealed 6 different cell states^41^. A recently reported single cell-resolved atlas of combined human breast cancers inferred 6 distinct clusters of neoplastic and malignant epithelial cells within in the TNBC subset^15^. While we do not know the relationship between the 6 different subpopulations identified by those orthogonal approaches, we assumed we would need to capture at least 6 different cell types from a given TNBC parental cell line. Assuming equal frequency, we calculated a 16.6% probability of capturing a given subpopulation, thus providing an expected yield of 16 wells of each subpopulation in a 96 well plate.

The number of resulting NBCs and BCs derived from any given cell line depends on the ability of the single cells to proliferate and form a colony; not all single cells survive the subcloning process. We also discarded wells in which we visualized more than one cell per well after the cell sorting/dilution plating step. Moreover, we accounted for the fact that the frequency of each of the 6 inferred cell states may not be equal. For these reasons, we standardly sort a minimum of 288 single cells (3 × 96 well plates) to achieve 30-40 different NBCs per cell line, and for each cell line, we mixed an equal number of cells from each BC to create BC Pools (Table S1).

To confirm that SunCatcher captures population complexity, we monitored cell morphology during the subcloning process. For example, the 31 Met1 BCs range across a spectrum from epithelial to mesenchymal phenotype; some of the individual BC clones have uniform morphology (e.g., BC26 is mesenchymal; BC45 is epithelial) while others appear heterogeneous (e.g. BC73) (Fig. 2A). We also confirmed that the Met1 BC Pool, which was comprised after mixing an equal number of each BC, is morphologically heterogeneous (Fig. 2A). Likewise, the HMLER-HR BC populations were also morphologically diverse (Fig. S1). We evaluated each Met1 BC for important tumor phenotypes known to impact tumorigenesis, disease progression, and response to therapy^42^. We focused on markers of stratified epithelium (cytokeratins 8 and 14), epithelial-mesenchymal transition (Zeb1 and EpCAM), and immune regulation (MHC-I and PD-L1).

**Figure 2.**
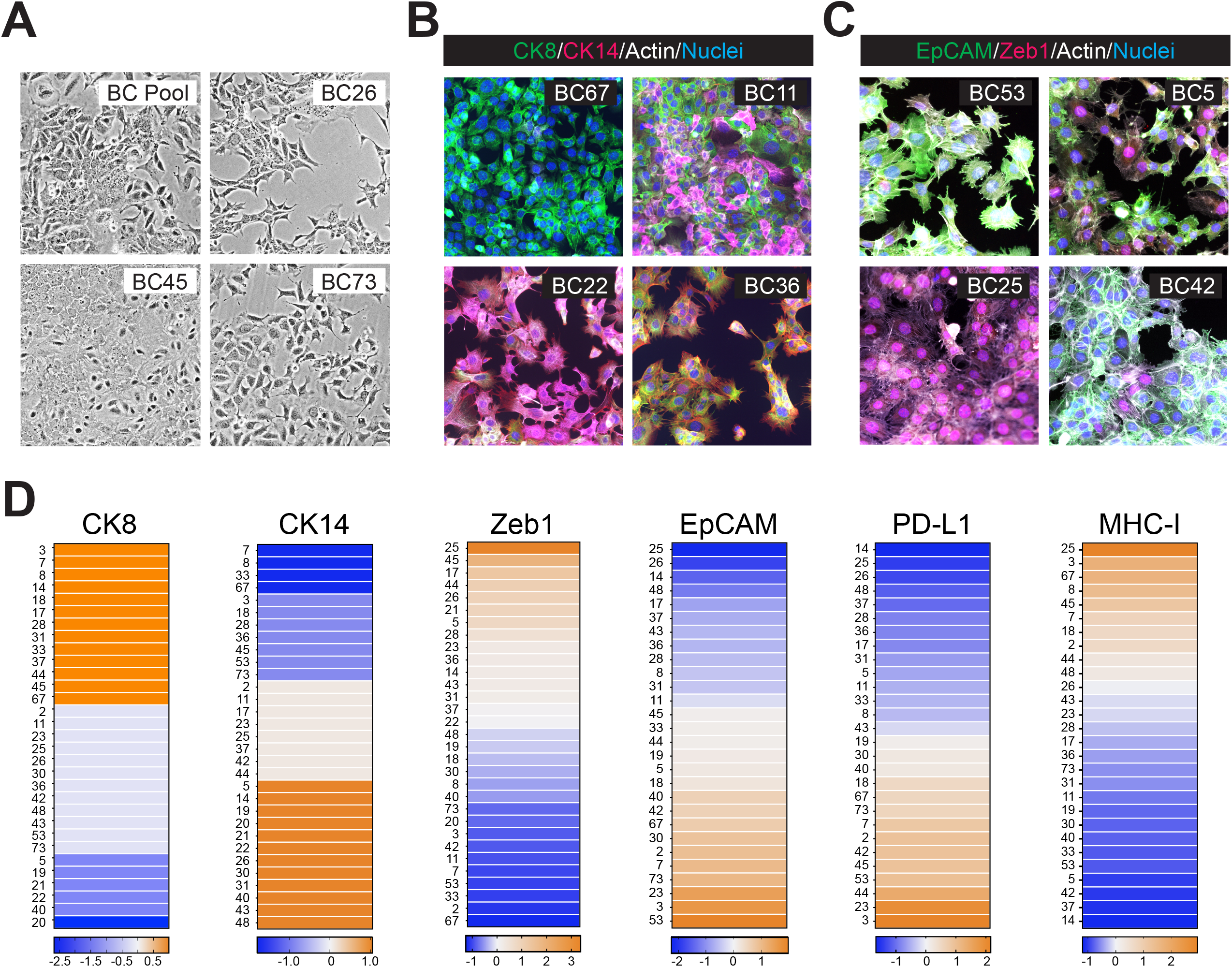
Phenotypic Diversity of Met1 Barcoded Clonal Populations. **(A)** Phase contrast microscopic images of indicated Met1 BCs; 20x magnification. **(B)** Immunofluorescence images of indicated BCs stained for cytokeratin 8 (CK8, green), cytokeratin 14 (CK14, red), f-actin (white), and nuclear counterstain (DAPI, blue); 20x magnification. **(C)** Immunofluorescence images of indicated BCs stained for epithelial cell adhesion molecule (EpCAM, green), Zeb1 transcription factor (red), f-actin (white), and nuclear counterstain (DAPI, blue); 20x magnification. **(D)** Relative expression levels of indicated factors across the Met1 BCs. CK8, CK14, and nuclear Zeb 1 were quantified by immunofluorescence image analysis; EpCAM, PD-L1, and MHC-I were assessed by flow cytometry and ranked by mean fluorescence intensity.

Immunocytochemical analysis of cytokeratin 8 (CK8, “luminal”) and cytokeratin 14 (CK14, “basal”) staining revealed that some BCs expressed a single cytokeratin (e.g., BC67 and BC22) (Fig. 2B). Other BCs contained cells that stained for either CK8 or CK14 to varying extents (e.g., BC11), and some BCs (e.g., BC36) contained individual cells that expressed both cytokeratins (Fig. 2B). Hence, despite their clonal origin, some BCs gave rise to phenotypically diverse populations. We quantified the cells expressing either marker as a percentage of all cells in each image and ranked BCs by their relative expression of each cytokeratin (Fig. 2D).

Similarly, co-staining for EpCAM and Zeb1 revealed that a few BCs almost exclusively expressed either nuclear Zeb1 (e.g., BC25) or EpCAM (e.g., BC53) (Fig. 2C. Most BCs (e.g., BC5 and BC42) expressed both factors to varying extents (Fig. 2C). We quantified the number of cells per field that expressed nuclear Zeb1 for each BC and ranked the BCs accordingly (Fig. 2D). Flow cytometry provides a more accurate quantification of EpCAM expression; therefore, we assessed cell-surface expression and ranked BCs by their mean fluorescence intensity (MFI) (Fig. 2D). We also evaluated cell-surface expression of PD-L1 and MHC-I for each BC by flow cytometry and ranked BCs according to the MFI for each factor (Fig. 2D).

### Next-Generation Sequencing-Based Barcode Detection and Quantification

The PRISM platform is useful for barcode deconvolution and has yielded important insights about susceptibility of various cancer cell lines to treatment and disease progression ^28, 31^. We sought to develop a next-generation sequencing (NGS)-based method of barcode detection in cases where multiplexing is desired. To do so, we first confirmed BC detection by Sanger sequencing using primers against the lentiviral barcode vector further upstream and downstream than those used for PCR amplification prior to PRISM-based barcode detection (Fig. S2A). Additionally, we used a DNA extraction method (Qiagen QiaAmp DNA Mini Kit) that yielded DNA samples that were less prone to degradation. Using this optimized DNA extraction and PCR amplification protocol, we validated that all 30 distinct barcodes could be amplified and detected from the HMLER-HR BCs using Sanger-sequencing (Fig. S2B, C).

In order to maximize the number of samples that we could sequence per Illumina sequencer lane, we chose to use multiple levels of multiplexing within our Illumina library preparation strategy. Twenty-four uniquely indexed Illumina adaptors are commercially available, enabling us to pool up to 24 samples within a single sequencing lane. Systems that allow for further multiplexing through addition of a second index during PCR amplification have been previously described^43^. We chose to introduce a second PCR step into our library preparation work-flow during which one of 20 unique indexes would be added to the 3’ end of the barcode amplicon (Fig. 3A, Table S2). Following PCR purification of the indexed barcode amplicons, we pooled equal molar amounts of each amplicon (Fig. 3A). Each barcode-index pool is then inserted into an indexed Illumina adaptor via a ligation reaction (Fig. 3A). Up to 24 unique barcode-index pools can be paired with the indexed Illumina adaptors, thereby allowing for the sequencing and deconvolution of up to 480 unique samples (20 barcode-index pairs x 24 indexed Illumina adaptors) in a single sequencing lane. Identification of the barcodes present within each sample thus requires deconvolution by first isolating all barcode reads associated with a single Illumina index; and second, by isolating all barcode reads within a single Illumina index that are associated with the specific barcode-index that is associated with that sample.

**Figure 3.**
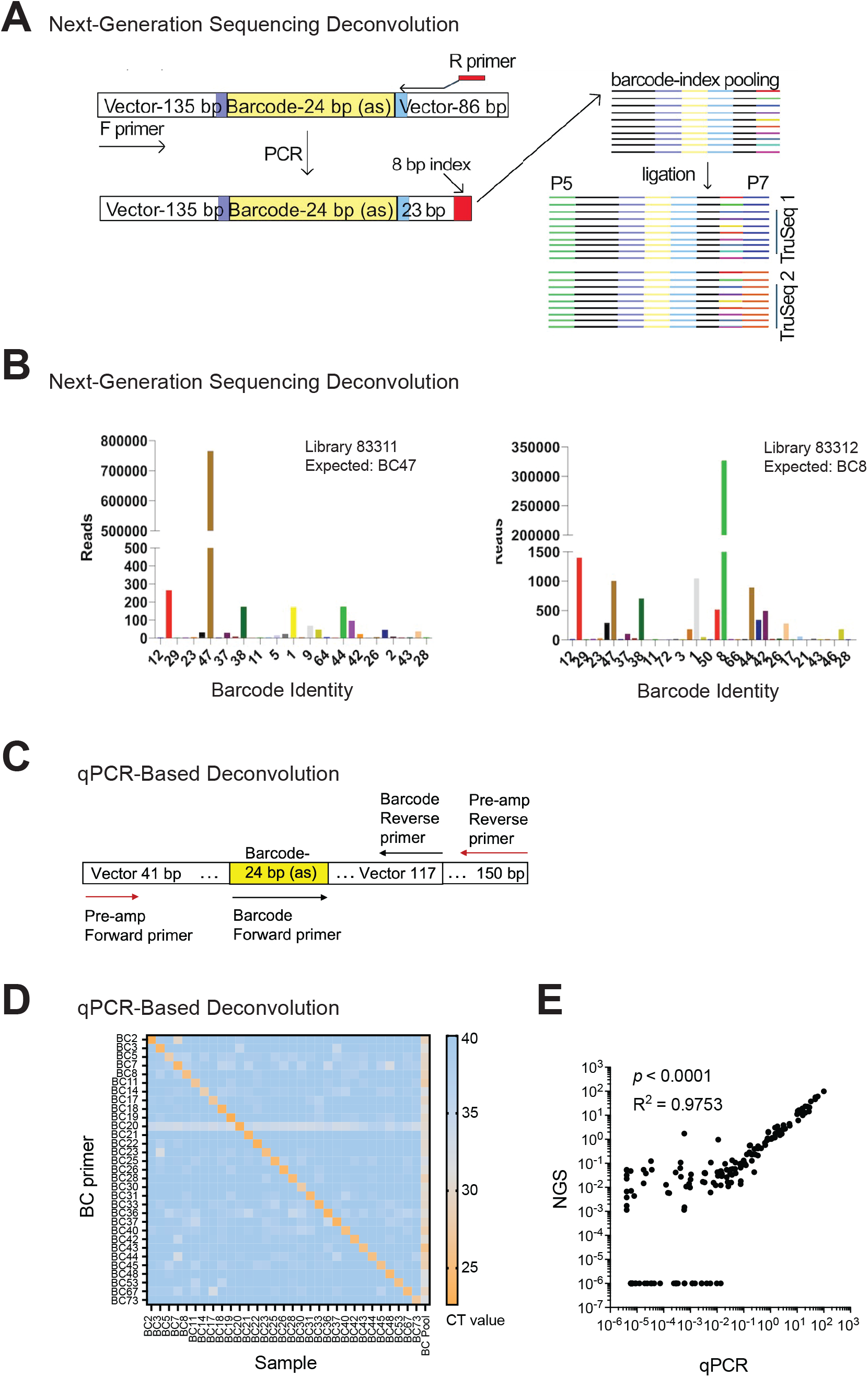
BC Detection by Next-Generation Sequencing and qPCR-Based Methods. (**A**) Multi-level multiplexing is achieved by attaching 1 of 20 unique barcode-indexes to the 3’ end of each sample. Up to 20 indexed samples can be pooled into a single barcode-index library. Each barcode-index library can be ligated into 1 of 24 uniquely indexed Illumina adaptors. Purple region represents PstI, light blue represents MluI. (**B**) Sequencing read counts for 2 test samples using the Illumina library preparation method. Each library corresponds to a single Illumina adaptor, and the expected barcode pair for each library is indicated. (**C**) Design of pre-amplification primers (red arrows) and BC-specific primers (black arrows). (**D**) Heatmap of qPCR C_T_ values when testing each indicated barcode oligonucleotide primer against every Met1 BC population as well as the Met1-BC pool. (**E**) Genomic DNA was isolated from 6 tumors derived from the McNeu BC Pool after ∼5 weeks of tumor growth in vivo; each barcode was quantified as a percentage of total barcodes in a given tumor by both qPCR and next-generation sequencing. Each datapoint represents the value for an individual barcode from an individual tumor by each method.

We tested each indexed primer for its ability to successfully amplify the barcode region of gDNA extracted from HMLER-HR BCs, and then used Sanger sequencing to verify that the index had been incorporated. All 20 indexes could be added to the 3’ end of barcode amplicons using PCR. We then tested whether our library preparation method could be used to accurately detect and deconvolute known barcode-index pairs that had been ligated into indexed Illumina adaptors. For this, we used PCR to pair each of 17 indexes with 2 different barcode sequences to generate 17 barcode-index pools. We used input DNA from single BCs, so that a single known DNA barcode would be present in each sample. Additionally, we indexed DNA samples from the HMLER-HR BC Pool (which contains 30 DNA barcodes) with 3 separate indexes to interrogate a situation where a larger number of barcodes are associated with each index.

These 20 samples were submitted to the Harvard Biopolymers facility (BPF) for Illumina library preparation. Each of the 20 barcode-index pools was ligated to a uniquely indexed Illumina adaptor. These adaptors have identical P5 regions, and their P7 regions each contain a unique TruSeq Index (Fig. 3A). These 20 multiplexed samples were then mixed and run on a single lane of an Illumina MiSeq sequencer. In order to adjust for the low sequence diversity present in the sample, 40% PhiX was spiked into the sequencing reaction to increase the sample diversity^44^.

If ligation efficiency is low, one cannot use PCR to amplify the ligation products using primers against the Illumina P5/P7 adaptor sequences because every barcode in the pooled sample would get equally amplified. Therefore, as an alternative to ligation-based Illumina library preparation methods, we attached the Illumina adaptor regions to the amplicons being sequenced using a PCR-based library preparation method. To pursue this method, we designed a single forward PCR primer with complementarity to the barcode vector and 20 reverse primers that each contained a unique Illumina TruSeq index (Table S3).

We generated a test library that contained five DNA samples from individual HMLER-HR BCs in order to evaluate whether this method enabled correct identification of the expected barcode-Illumina index pairs. The PCR reactions for each primer set were highly specific and efficient, generating a single strong band of the expected length (Fig. 3B, S3B). Importantly, the expected barcode-Illumina index pairs generated signal that was ∼233-2,500-fold higher than the highest false positive signal, allowing for simple thresholding of true signal from background sequencer noise (Fig. 3B, S3B). Together, these data demonstrated that the barcodes were amenable to next-generation sequencing-based detection.

### qPCR-Based Barcode Detection and Quantification

We reasoned that qPCR would be a rapid, inexpensive method to identify and quantify barcodes from our BC Pools owing to the smaller library size of SunCatcher compared to traditional barcode libraries, which typically contain 10^6^-10^8^ barcodes. We first designed oligonucleotide primers to common barcode flanking sequences to enable a pre-amplification step (Fig. 3C). The pre-amplification step has several advantages. First, a separate PCR reaction is required to detect each barcode, so pre-amplification would ensure sufficient gDNA input material, particularly from precious samples. Second, it adds a layer of specificity for BC detection above any potential background signal from the non-barcoded genome. Finally, it enriches all barcodes, including any very low abundance barcodes so that their relative contribution to any sample can be quantified. We also designed oligonucleotides specific to each barcode sequence (Fig. 3C, Table S4).

To ensure the specificity of our PCR detection method, we tested each individual barcode oligonucleotide primer against each individual BC population as well as the BC Pool from the Met1 murine triple-negative breast cancer cell line and plotted the C_t_ values for each reaction. Every barcode was detected in the BC pool, while each oligonucleotide primer amplified only its specific barcode (Figure 3D). Note that if any of the primers had amplified a non-specific barcode, we would have been able to retrieve the corresponding NBC population and infect it with a different barcode until we achieved specificity, thus demonstrating another advantage to the SunCatcher subcloning approach.

We directly compared results using NGS to those using qPCR on the same tumors derived from the McNeu BC Pool (n=6). We found significant concordance and correlation in barcode composition between methods (R^2^ = 0.975; p<0.0001; Figure 3E). The qPCR-based detection method proved more sensitive than NGS for low-abundance barcodes, given that NGS requires thresholding to distinguish between low-level false positive signals and signal generated by true, low-frequency barcode variants (Fig. 3E).

Hence, barcodes can be accurately detected by both NGS and qPCR methods. Our NGS protocol enables multiplexing for simultaneous generation of large amounts of barcode information, particularly if one wishes to use larger barcode libraries. Our qPCR protocol is much more rapid, inexpensive, and highly sensitive for smaller barcode libraries.

### SunCatcher Barcoding Retains Biological Properties of Parental Tumor Cells

Confirming that BCs captured heterogeneity of the parental cell lines and that our PCR-based detection method was specific and sensitive, we mixed an equal number of each BC to create BC Pools (Table S1). We first evaluated BC Pool compositions in culture. The Met1 BC Pool composition shifted with time in culture; while some clones expanded (e.g., BC45), no clones dropped below the limit of detection (Fig. 4A). The HMLER-HR BC Pool was more stable in culture (Fig. S4A). Importantly, the growth rates of the Met1 BC Pool (Fig. 4B) and the HMLER-HR BC Pool (Fig S4B) were nearly identical to those of their respective parental cell lines.

**Figure 4.**
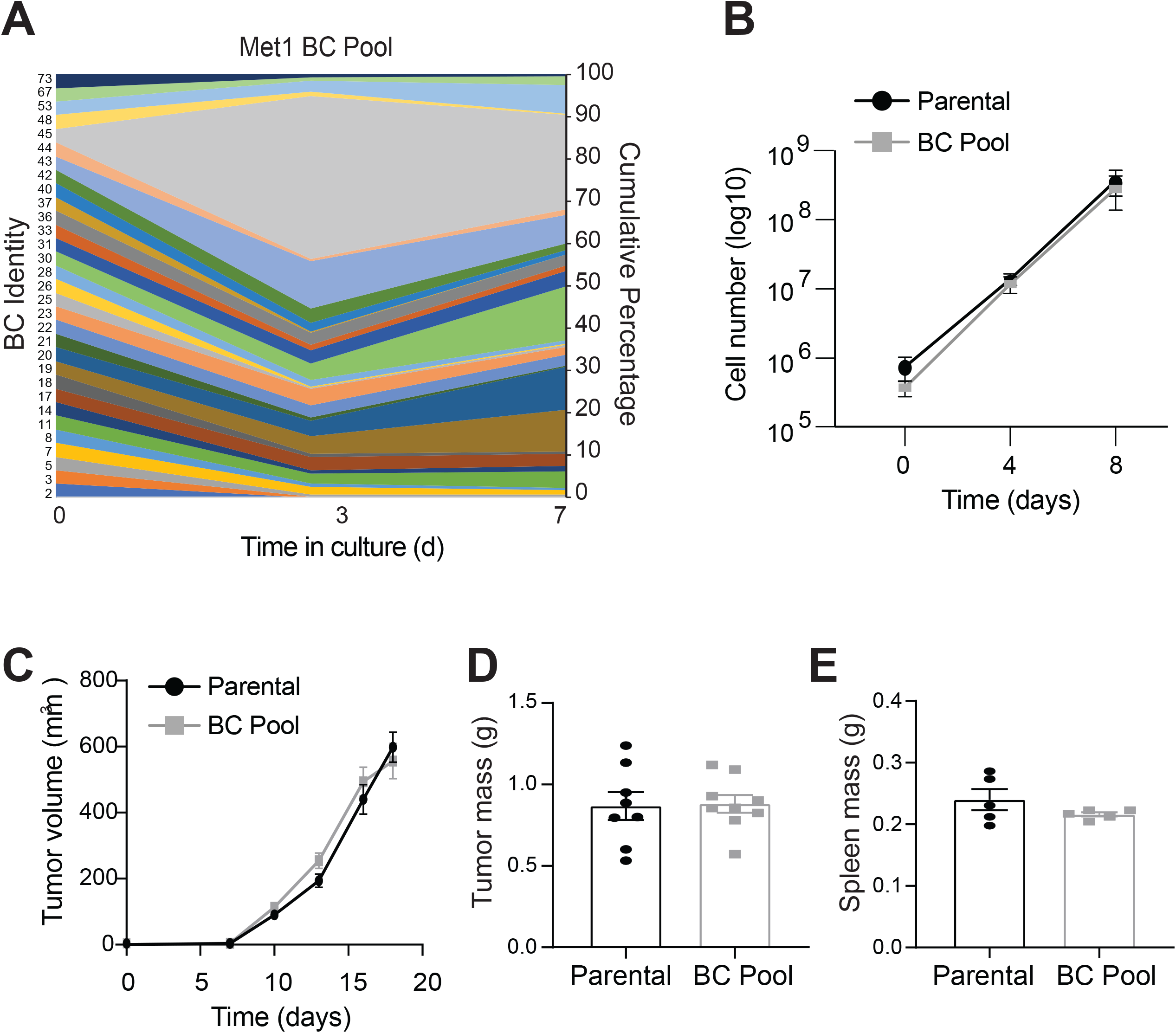
SunCatcher Barcoding Retains Tumorigenic Properties of Parental Cell Line. **(A)** Sand plot showing clonal composition (cumulative percentage) of Met1 BC Pool over 2 passages *in vitro*. **(B)** Growth kinetics of Met1 Parental (black; Y = 0.3360x + 0.1699) and Met1 BC Pool (grey; Y = 0.3562x + 0.4121) over 8 days in culture; n=4 per group; error bars = SD. **(C)** Growth of tumors from Met1 parental cells and Met1 BC Pool cells in FVB mice; n=5 per cohort, error bars represent S.E.M. **(D, E)** Mass (g) of tumors (D) and spleens (E) from experiment represented in C.

It was most important for us to learn whether the SunCatcher “deconstruction-reconstruction” approach affected in vivo tumor biology; therefore, we orthotopically injected either 2.5×10^5^ Met1 Parental cells or 2.5×10^5^ Met1 BC Pool cells into cohorts of FVB mice (n=9 per cohort), using cells after only 1 passage in culture. There were no significant differences between cohorts in tumor growth kinetics during the experimental time course or in the final mass of tumors or spleens at the experimental end point (Fig. 4C-E). Likewise, we observed no significant difference in tumor growth or final tumor mass between the parental 4T1 cells and the 4T1 BC Pool (Fig. S4C).

Collectively, these data revealed that the SunCatcher clonal barcoding approach preserved the heterogeneity as well as in vitro and in vivo growth kinetics of the original tumor cell lines from which they were derived. Moreover, subpopulations of just ∼30 clones were sufficient to recapitulate phenotypic and functional properties of their respective parental populations.

### Using SunCatcher to Understand Dynamics of Primary Tumor Growth

We asked how the BC composition compared between individual tumors and if we could devise a method for representing average BC composition across individual tumors of a given cohort. To do so, we injected 2.5×10^5^ Met1 BC Pool cells orthotopically into contralateral mammary fat pads of FVB mice (n=5). As a control, we analyzed a sample from the injection material that was taken at the time the cells were prepared for injection (“pre-injection” sample) and a sample taken after the last mouse was injected (“post-injection” sample) to determine if the BC composition shifted during the time (∼2 hr) it took to complete all injections. The composition of the pre-and post-injection samples were identical (Table S6); hence, all mice in the cohort received the same input (Fig. 5A).

**Figure 5.**
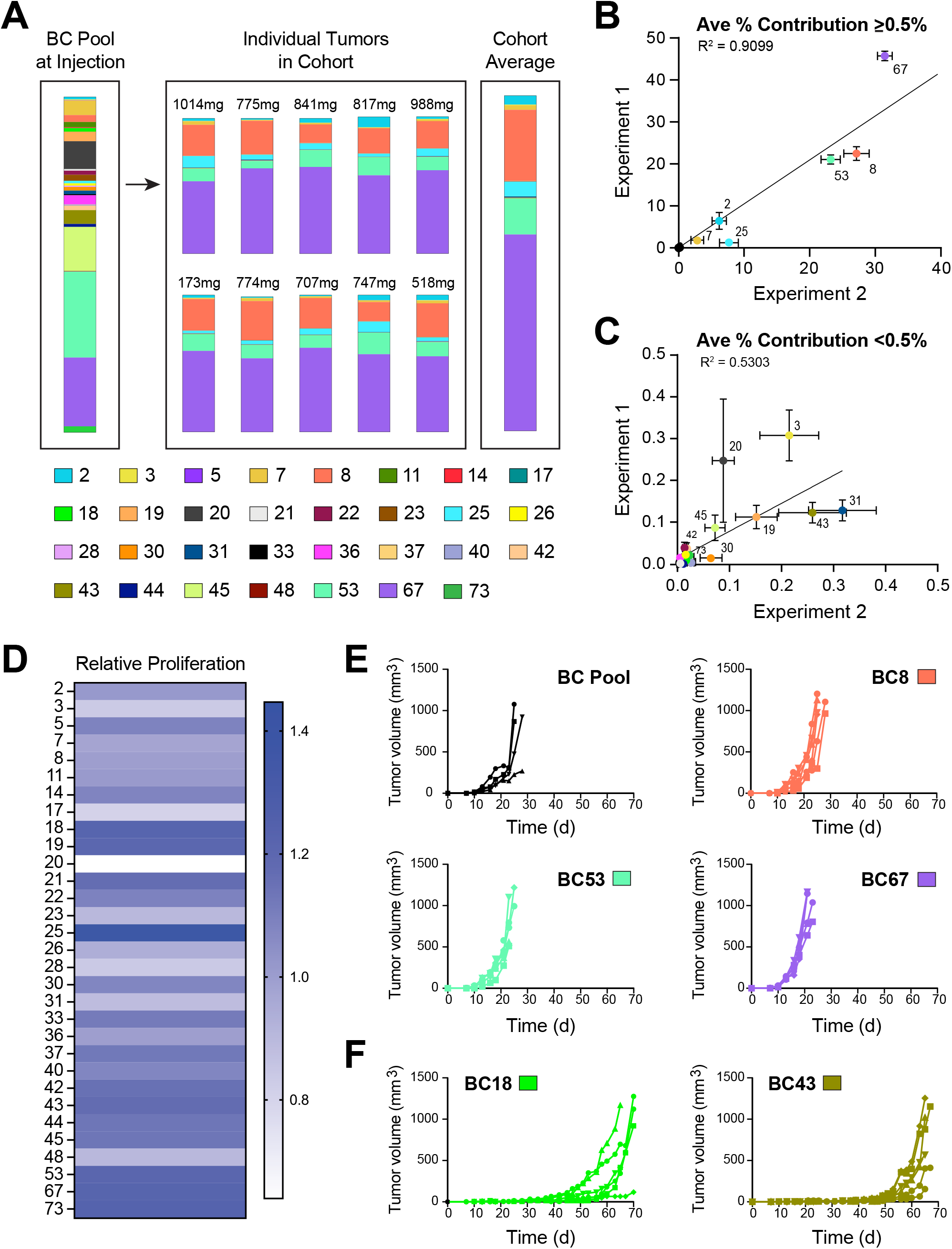
SunCatcher Enables Analysis of Clonal Dynamics During Primary Tumor Progression. **(A)** Quantitative PCR assessment of barcode composition in the Met1 BC Pool at time of injection (left), in each of 10 tumors (n=5 mice) after 18 days (center), and calculation of the average composition of all tumors (right). Composition bars show indicated barcodes as a percent of total barcode signal (100%) within each sample; tumor mass (mg) is indicated above each bar. Color code for each BC is indicated. **(B, C)** Average representation of each BC in n=10 tumors for each of 2 experiments. BCs that constituted >0.5% of total BC signal are represented in (B) and <0.5% are shown in (C). The correlation coefficients (R^2^, simple linear regression) are shown; error bars represent S.E.M. **(D)** Heatmap of relative proliferation rate (doubling time) of each individual BC in culture over a 9-day period. **(E)** Growth of tumors derived from the Met1 BC Pool or indicated individual BCs. 2.5 × 10^5^ cells were injected orthotopically into FVB mice (n=5 per cohort) and allowed to grow until ethical end point of ∼1000 mm^3^. These BCs represent the 3 most dominant clones from experiments in A-C. **(F)** Growth of tumors derived from the indicated BCs. 2.5 × 10^5^ cells were injected orthotopically into FVB mice (n=6 per cohort) and allowed to grow until ethical end point of ∼1000 mm^3^. These BCs represent 2 of the BCs that comprised <0.3% of the tumors in experiments from A-C.

After a growth period of 18 days, we harvested the tumors, isolated genomic DNA, and used our PCR-based method to quantify the contribution of each barcode to each tumor. We observed remarkable similarity in BC composition across all 10 tumors in the cohort despite differences in tumor mass (Fig. 5A, Table 1), suggesting that tumor-forming capacity of this pre-clinical TNBC model is not stochastic. Given the consistency between mice, we were able to represent the average BC composition for the entire cohort (Fig. 5A).

**Table 1.** BC Composition of Orthotopic Primary Tumors. (Related to data shown in Fig. 5A). The percent composition of each BC in the Met1 BC Pool at the time of injection is represented as “Pre-Injection”. The average contribution of each BC as a percent of total BCs in each resulting tumor (n=10) and S.D. are indicated in respective columns.

|  | % Composition |  |  |
| --- | --- | --- | --- |
| BC | Pre-Injection | Tumor Average | Tumor SD |
| 2 | 0.7459 | 2.6848 | 2.0588 |
| 3 | 0.2218 | 0.2247 | 0.1327 |
| 5 | 0.1089 | 0.0058 | 0.0034 |
| 7 | 4.3510 | 1.3084 | 0.6790 |
| 8 | 2.0571 | 21.2297 | 4.8215 |
| 11 | 1.5696 | 0.0050 | 0.0057 |
| 14 | 0.2405 | 0.0002 | 0.0002 |
| 17 | 0.0207 | 0.0003 | 0.0007 |
| 18 | 1.0353 | 0.0018 | 0.0035 |
| 19 | 2.8757 | 0.1045 | 0.0507 |
| 20 | 8.3150 | 0.0693 | 0.0357 |
| 21 | 0.4561 | 0.0003 | 0.0005 |
| 22 | 1.2243 | 0.0108 | 0.0111 |
| 23 | 1.7757 | 0.0417 | 0.0353 |
| 25 | 0.7883 | 4.3674 | 2.2510 |
| 26 | 0.6720 | 0.0011 | 0.0020 |
| 28 | 0.3914 | 0.0010 | 0.0024 |
| 30 | 1.0937 | 0.0070 | 0.0083 |
| 31 | 1.0464 | 0.2874 | 0.1328 |
| 33 | 0.3400 | 0.0154 | 0.0187 |
| 36 | 2.7291 | 0.0404 | 0.0204 |
| 37 | 0.0587 | 0.0019 | 0.0043 |
| 40 | 0.2769 | 0.0198 | 0.0075 |
| 42 | 1.3537 | 0.0130 | 0.0149 |
| 43 | 4.1184 | 0.1688 | 0.0679 |
| 44 | 0.8892 | 0.0060 | 0.0073 |
| 45 | 13.1404 | 0.0603 | 0.0349 |
| 48 | 0.1662 | 0.0067 | 0.0069 |
| 53 | 25.6185 | 10.7190 | 2.8135 |
| 67 | 20.5118 | 58.5972 | 3.7022 |
| 73 | 1.8079 | 0.0001 | 0.0003 |

We repeated the orthotopic tumor growth experiment to determine reproducibility of results between experiments. For BCs that constituted >0.5% of the total tumor composition, we observed concordance between results from the 2 different experiments (R^2^=0.9099; Fig. 5B). There was more variability within and between experiments for those BCs that represented <0.5% of the total tumor BC composition (R^2^=0.5303; Fig 5C). The reproducibility of our results suggested phenotypic stability among the clones.

It has been hypothesized that proliferation rates dictate clonal dynamics and that the most proliferative clones will dominate a tumor^4, 45^. However, what provides the greatest selective pressure during disease progression has not been well elucidated. For example, it is not clear whether the most dominant clones in a tumor are inherently more proliferative or whether their emergence is a selective process. Therefore, we tested individual Met1 BCs *in vitro* and *in vivo* to understand how their growth properties related to their fate within the resulting tumors. Three clones together consistently comprised ∼90% of total BCs detected in the tumors at the experimental end point: BC8 (21.2 ± 4.8%), BC53 (10.7 ± 2.8%), and BC67 (58.6 ± 3.7%) (Fig. 5A-C; Table 1). BC53 and BC67 were among the most dominant clones in the BC Pool at the time of injection (25.6% and 20.5%, respectively); hence, the contribution of BC53 contracted ∼58% while BC67 expanded ∼185% *in vivo* (Table 1). These two BCs had similar proliferation rates *in vitro* (Fig. 5D), and they each displayed similar latency and rapid growth kinetics *in vivo* when injected as pure clonal populations (Fig. 5E). In contrast, BC8 comprised only ∼2% of the BC Pool at the time of injection (Table 1) and was among the slowest proliferators *in vitro* (Fig. 5D). Nevertheless, the relative contribution of BC8 expanded ∼10.5-fold *in vivo* (Table 1) and it displayed relatively short latency and rapid growth kinetics when injected as a pure population (Fig. 5E). Those results suggested that BC8 experienced positive selective pressure *in vivo*, causing it to become one of the most rapidly proliferating clones.

Other clones appeared to undergo negative selection *in vivo*. For example, BC45 represented a relatively large portion (∼13%) of the BC Pool at time of injection and was highly proliferative *in vitro*, yet it represented only ∼0.06% of the resulting tumors (Fig. 5A-E; Table 1). Other clones, such as BC14, BC17, BC21, and BC73, diminished to only <0.0004% of the tumors (Table 1).

Even more strikingly, BC18 was one of the most proliferative clones *in vitro* and yet it comprised only ∼1% of the BC Pool at the time of injection and comprised <0.002% of the resulting BC Pool tumors (Fig. 5D; Table 1). Likewise, BC43 comprised ∼4% of the BC Pool at time of injection and was diminished to only <0.2% of the resulting tumors (Table 1). Nevertheless, BC18 and BC43 were indeed capable of forming tumors on their own *in vivo*, albeit with long latency (∼40-50 days) and variable incidence (Fig. 5F). Those results lead one to speculate that such minor clones might persist and achieve their tumor forming capacity should they be given sufficient time to do so.

Collectively, these results suggested that clonal dynamics and/or environmentally driven selection pressures ultimately influence the fate of certain BCs while the inherent proliferation of other BCs is not impacted. Although these concepts have been well established, SunCatcher enabled us to identify the specific clones within heterogeneous tumors that are influenced by such dynamics *in vitro* and *in vivo*.

### Using Suncatcher to Study Metastasis

We typically do not detect Met1 spontaneous metastasis prior to 30 days using conventional detection methods. Historically, we have only detected metastases in the lungs of a portion of any given cohort before mice reach a humane end point due to primary tumor burden. Given the sensitivity of barcode detection and reproducibility of results, we asked whether we might detect early Met1 BC Pool metastases from orthotopic sites. To do so, we injected the Met1 BC Pool into contralateral mammary glands and examined visceral and skeletal tissues 3 weeks later. Mice were otherwise healthy and we did not observe any visible, overt metastases in any tissues. However, barcode detection indicated that the lungs, long bones, and mandibles consistently contained barcode signal.

We also devised a method by which we would be able to extrapolate numbers of metastatic cells based on total barcode qPCR signal obtained from a fixed mass of tissue (see Methods). Our method enabled thresholding to distinguish real signal from background for each tissue (see Methods and Fig. 6A). Consequently, we were able to reproducibly detect <1 cell per 0.1 mg of tissue (Fig. 6A).

**Figure 6.**
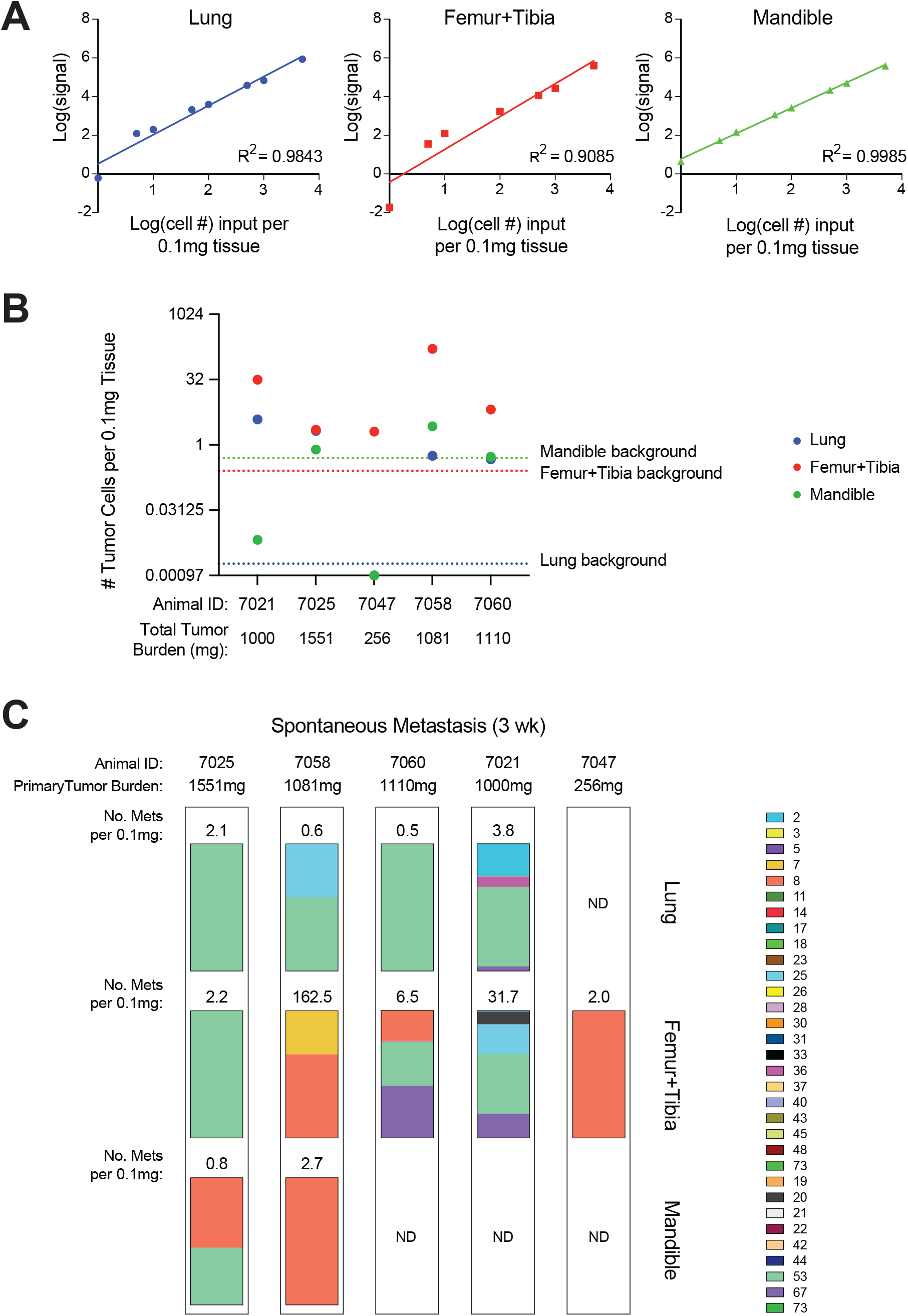
SunCatcher Barcoding to Study Metastasis. **(A)** Calibration were curves generated for indicated tissues by serially diluting known amounts of barcoded tumor cell gDNA into a fixed amount of normal tissue gDNA. From top to bottom: lung, long bones (from femur and tibia), mandible. **(B)** 2.5×10^5 barcoded Met1 tumor cells were injected bilaterally into the mammary fat pads (n=5 animals) and tumors were allowed to grow for 21 days, at which point tissues were harvested and metastasis burden was calculated. Dashed lines indicate the background signals from each indicated tissue type. Tissues with signal above the background were considered as positive for metastasis and estimated tumor cell number per 0.1mg tissue was calculated based on the calibration curve for that specific tissue. **(C)** Barcode composition analysis on tissues with positive metastasis signal. Bars represent percent of total barcode signal (100%) within each sample. Also shown are mouse identities, total primary tumor burden for each animal, and estimated numbers of metastases per tissue; N.D., not detected.

Total primary tumor burden was similar between 4/5 mice (mean 1185.5mg; range 1000-1551mg) and only one small tumor (256mg) formed in one mouse (Fig. 6B). Among the tissues in which we detected barcode signal, the leg bones (femur and tibia combined) had the greatest relative numbers of spontaneous metastases, regardless of primary tumor burden, whereby 100% of the mice had barcode signal above threshold (Fig. 6B). Lung metastasis was detected in 4/5 mice and metastasis to the mandible was detected in 2/5 mice (Fig. 6B).

BC53 was a dominant clone in 100% of mice that had lung metastasis and the only clone detected in 50% of those mice (Fig. 6C). In the leg bones (femur and tibia), the inter-individual composition of BC metastases was slightly more variable and diverse. For example, unlike in lung, BC53 was detected in 60% of the mice with metastases to the leg bones (Fig. 6C). The mice with the lowest tumor burden (mouse 7047; 256 mg) and highest tumor burden (mouse 7025; 1551 mg) each had clonal leg bone metastases (BC8 and BC53, respectively; Fig. 6C). BC8 and BC53 were the only barcodes detected in the mandibles, and here, BC8 was the dominant clone (Fig. 6C).

We were also able to assess intra-individual differences in composition across different metastatic sites. Mouse 7025 displayed relative consistency across all tissues, with BC53 clonal metastasis in lung and leg bones and BC8 appearing only in the mandible (Fig. 6C). Interestingly, mouse 7047, which had the lowest primary tumor burden, bore BC8 clonal metastasis to the leg bones (Fig. 6C). Those results suggested that BC8 is particularly amenable to bone metastasis.

Collectively, these results indicated that SunCatcher offers a functional barcoding approach to study clonal dynamics during metastatic progression. Importantly, the qPCR-based detection method enabled us to identify and quantify metastases even at very early stages.

## DISCUSSION

A number of innovative barcoding technologies have been developed that enable one to study clonal dynamics among heterogeneous populations of cells across a variety of applications^19-34^. Many original barcoding methods rely on infecting heterogenous populations of cells with libraries of 10^6^-10^8^ barcodes without the ability to isolate and study every individual clone. Due to the single cell cloning approach, SunCatcher enables functional analysis of individual barcoded populations and generation of custom pools of populations of cells of interest. Each clone is specific to a single barcode; there is no chance that a clone contains more than one unique barcode sequence and thus, will not confound analyses or data interpretation. Moreover, our PCR-based BC detection method is inexpensive, rapid, sensitive, and reproducible and our NGS-based method allows for multiplexing for larger barcode libraries. While generation of individual clones is labor-intensive up front, once NBC and BC stocks are made, those stocks can be used indefinitely. We envision that SunCatcher could easily be adapted for use in any cell-based study and hope it will be useful to the research community.

We used reported data from analyses of normal breast epithelium and breast cancer to estimate the number of clones required to capture sufficient heterogeneity from the parental cell lines. Theoretically though, the only limit to the numbers of BCs that can be generated is the ability to manage many single cell-derived clones. The usefulness of the PCR-based detection method depends on barcode sequence design and oligonucleotide primers specificity. However, our next-generation sequencing protocol for BC detection should render the SunCatcher clonal system scalable to larger numbers.

Our analyses of BC cell phenotypes exposed additional benefits of the SunCatcher system. First, our clonal approach enables one to trace clonally related cells, even as the inherent plasticity of those cells gives rise to phenotypic heterogeneity. Second, one can also distinguish different clones from among groups of cells that share common phenotypes. Because individual clones are retained, SunCatcher enables one to not only identify but also isolate the common ancestor of cells that might otherwise appear to be different from one another by other analysis methods. While not performed here, the Suncatcher system will allow for unbiased transcriptional profiling or more specialized functional assays to further characterize the degree of functional heterogeneity among each clone in a barcoded pool.

As we applied SunCatcher to pre-clinical breast cancer models, we found that a BC Pool of just 31 clones was sufficient to recapitulate tumor growth kinetics of the parental population. On their own, the individual BCs had different growth rates *in vivo*, suggesting that heterogeneity was stable and that clonal dynamics contribute to the overall growth rate of a tumor that is comprised of phenotypically different clones.

Most often, end points in pre-clinical cancer research are dictated by humane tumor size. Using SunCatcher, we identified clones (e.g., BC43) that contributed <0.3% of the tumor BC composition at our end point *in vivo* yet had the potential to form tumors after a long latency period when injected as a pure clonal population. Such findings raise questions about the fate of those clones if our experimental time points were extended or if those clones represented residual disease after surgical resection or treatment. In fact, several studies provide evidence that metastases are derived from subclones present at low frequency in the primary tumor^46-48^. A better understanding of these minor clones could aid in the identification of therapeutic targets that could prevent disease progression.

In our experience, we had been unable to detect early metastases in our pre-clinical models using conventional detection methods. Moreover, we had assumed that our Met1 cells were unable to spontaneously metastasize to distant organs before primary tumors reached ethical end point. Our detection of spontaneous metastases in lungs and skeletal tissues after only 3 weeks are in line with numerous studies that have detected circulating and disseminated tumor cells in the early stages of cancer progression in both preclinical models and in patients^49-52^. We envision that SunCatcher could provide valuable information about the properties of individual disseminated tumor cells that make them more or less threatening than their counterparts.

Perhaps one of the more important benefits of SunCatcher is that it not only enables analysis of clones that persist during disease progression or that resist treatment, but also enables one to study clones that either do not form tumors or are responsive to treatment. The ability to study “super responders” could yield important information about their vulnerabilities and lead to therapeutic approaches to prevent disease progression.

## Methods

### Lentiviral Barcode Vectors

Barcode sequences were originally developed by Tm Bioscience (Toronto, Ontario, Canada)^53^ and detailed in the Luminex FlexMAP Microspheres Product Information Sheet^54^, as published by Peck, et al.^35^. Lentiviral vectors containing barcode sequences were generated as previously described^28, 35^. 293T cells were transfected with 2.5μg of lentiviral vector barcode DNA, 2.5μg pCMV-dR8.2 dvpr (Addgene plasmid 8455), 1μg pCMV-VSVG (Addgene plasmid 8454) in Opti-MEM Reduced Serum Medium (Gibco) with Fugene HD (Promega) or FuGENE6 (Roche Corporation). Viral supernatants were collected 48- and 72-hours following transfection and were filtered through a sterile 0.45μm syringe filter (VWR), and stored at −80°C.

### Cell lines

The Met1 TNBC cell line, which was originally generated from a spontaneously arising tumor in an FVB/N-Tg (MMTV-PyVmT) mouse^38^, was a gift from J. Joyce (University of Lausanne) with permission from A. Borowsky (UC Davis School of Medicine) and maintained in DMEM containing 10% fetal bovine serum, 100 U/mL Penicillin-Streptomycin, and 2mM Glutamine. The 4T1 TNBC cell line, which was derived from a spontaneously arising tumor in a Balb/c mouse^37^, was provided by F. Miller (Wayne State University School of Medicine) and maintained in DMEM containing 10% fetal bovine serum, 1x MEM Non-Essential Amino Acids Solution (Gibco™), 100 U/mL Penicillin-Streptomycin, and 2mM Glutamine. The McNeuA HER2+ cell line, which was originally derived from a spontaneously arising breast tumor in an MMTV-neu transgenic mouse, was provided by Michael Campbell (University of California, San Francisco) and maintained in DMEM (Gibco) supplemented with 10% fetal bovine serum (FBS) and 1% Penicillin Streptomycin (Gibco) at 37C with 5% CO_2_^36^. Human HMLER-hygro-H-*ras*V12 (HMLER-HR) cells^39, 55^ were cultured in advanced DMEM/F12 (Gibco) supplemented with 5% calf serum (CS, HyClone), 0.1% hydrocortisone (Sigma Aldrich), and 1% Penicillin Streptomycin (Gibco) at 37C with 5% CO_2_. 293T cells were cultured in DMEM (Gibco) supplemented with 10% FBS (Gibco) and 1% Penicillin Streptomycin (Gibco) at 37C with 5% CO_2_. All cell lines tested negative for mycoplasma (Lonza); short tandem repeat analysis verified the identity of mouse (Bioassay Methods Group, National Institute of Standards and Technology) and human (Promega GenePrint 10 System) cells.

### SunCatcher Barcoding Protocol

Single cells were isolated from cell lines by either FACS or plating 0.5 cell/well in 96-well plates. We verified that each well contained a single cell by phase microscopy under 40x magnification and discard wells that contained more than one cell. Each single cell clone was expanded and designated as non-barcoded clones (NBCs). Aliquots of 5 × 10^5^ cells of each NBC were prepared in appropriate culture medium + 10% DMSO and stored in cryovials in liquid nitrogen.

For barcoding, each NBC population was thawed into a 6-well plate and infected with a unique barcode-containing lentivirus supernatant in medium with Polybrene (Sigma Aldrich) at a concentration range from 1:200 to 1:50. After 24 hours, virus was washed out and replaced with fresh complete medium with 10μg/ml blasticidin (Sigma). After 3 days, cell numbers were counted in each condition and those with an infection rate of <10% (number of infected cells remaining in selection media / infected cells grown in regular growth media) were selected. Single cells were then isolated from the infected populations by plating 0.5 cell per well into 96-well plates and then expanded in order to generate multiple individual clonal populations for each barcode. From those, a single clonal population was selected at random to represent the barcoded clonal population (BC) for each unique barcode sequence. Aliquots of 5 × 10^5^ cells of each BC were prepared in appropriate culture medium + 10% DMSO and stored in cryovials in liquid nitrogen.

For each cell line, equal numbers of every BC were mixed to form a BC Pool. Aliquots of 5 × 10^5^ cells of each BC Pool were prepared in appropriate culture medium + 10% DMSO and stored in cryovials in liquid nitrogen.

### Cell proliferation Assay

Met1 parental or barcoded clones were plated to a density of 20,000 cells per well of a 6-well plate. Every 72 hours over a 9-day time course, cells were counted using a hemocytometer and replated to a density of 20,000 cells per well. Relative proliferation rates of each BC were represented as cell doubling time per day.

### Immunofluorescence and image analysis

Glass coverslips (#1.5; Election microscopy science) were coated with rat tail collagen I (Thermo-Fisher Scientific) overnight at room temperature. The coverslips were washed once with PBS and cells were seeded and spread overnight at 37°C and 5% CO_2_. The cells were then fixed in 4% Paraformaldehyde (Sigma), permeabilized in 1% TX-100 (Sigma) and blocked in 3% bovine serum albumin in PBS (Fisher Scientific). Cells were stained with the indicated primary antibodies for overnight in 4 °C, followed by 1 hour incubation for secondary antibodies in room temperature. Then nuclei were visualized with DAPI (1 μg/ml, Sigma) and mounted onto glass slides. Stains and primary antibodies: F-Actin (Rhodamine Phalloidin, 1:1000; Thermo Fisher,), CK14 (1:200; Biolegend), CK8 (1:10; TROMA-1, DSHB). Secondary antibody (Invitrogen, 1:250): anti-rat 647, anti-rabbit 488, anti-rat 488, anti-rabbit 647. Tissue slides were imaged using a Nikon Eclipse Ni microscope. CK8 and CK14 staining was quantified by two researchers on a 1-4 expression level scale where 1= 0-5%, 2= 6-30%, 3= 31-60% and 4=61-100% from 5 randomly selected images for each barcoded cell line.

### Flow cytometry

10^5^ cells were harvested using 0.25% Trypsin, centrifuged, and seeded into round bottom 96-well plates. The cells were washed with FACS buffer (PBS with 2% FBS) and centrifuged at 300 x g for 5 minutes at 4°C. Cells were incubated with anti-EpCAM (Clone G8.8, APC-Cy7, 1:400 dilution, BioLegend), anti-MHC-I (clone: KH114, FITC, 1:400 dilution, BioLegend), and PD-L1 (clone: 10.F.9G2, PE-Cy7, 1:100 dilution BioLegend) on ice for 30 minutes. The cells were washed twice with FACS buffer and analyzed on a BD Canto II flow cytometer. Data were analyzed using FlowJo software.

### Animal experiments

All animal studies were conducted in accordance with regulations of the Institutional Animal Care and Use Committee of the Brigham and Women’s Hospital (protocol no. 2017N000056). Female FVB mice 7 weeks of age were purchased from The Jackson Laboratory. Mice were 8-9 weeks of age at the time of injection. For orthotopic injections, 2.5×10^5^ Met1 (or Met1BC) cells were prepared in 20μl PBS and injected into the inguinal mammary fat pads. Tumors were measured with calipers 2-3 times per week, and tumor

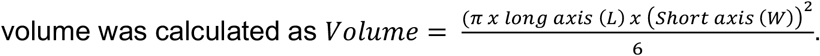

For spontaneous metastasis studies, 2.5×10^5^ Met1 BC Pool cells were prepared in 20μl PBS and injected into contralateral inguinal mammary fat pads of FVB mice. Mouse weight was monitored daily, and animals were sacrificed if their weight dropped by 20%. Otherwise, mice were sacrificed 21 days after injection by CO2 inhalation, perfused with PBS, and tissues were collected for barcode analysis.

### Genomic DNA Preparation from Cells and Tissues

Cultured cells were detached using 0.25% trypsin, pelleted, resuspended in Buffer AL (Qiagen), and genomic DNA (gDNA) extracted using the QiaAmp DNA Mini Kit (Qiagen) according to manufacturer’s instructions. Tumor and other tissues were flash frozen in liquid nitrogen and pulverized using a mortar and pestle. 25mg of tissue was used for DNA isolation according to the manufacturer’s protocol (Qiagen). Femurs and mandibles were immediately dissected and cleaned to remove muscle and connective tissue and then flash frozen and pulverized (preparations included marrow). 25mg of bone preparations were used for DNA isolation according to the manufacturer’s protocol (Qiagen). DNA concentration was quantified using a Nanodrop 8000 Spectrophotometer (Thermo Fisher).

### Processing gDNA and next-generation sequencing

From genomic DNA preparations, the barcode region of the lentiviral barcode vector was amplified using either primer set JO primer F/R (F: 5’-TGGAGCATGCGCTTTAGCAG; R: ATCGTTTCAGACCCACCTCC-3’) or indexed primer sets (F: JH p05: 5’-ATC GTT TCA GAC CCA CCT CCC-3’; R: JH p11-14 and JH p31-46, Table S4). PCR reaction mixtures contained 250 ng DNA, OneTaq 2x MasterMix (New England Biolabs), and 10 μM each of F/R primers. The following PCR program was used to PCR amplify the template DNA: 94°C for 30 seconds; 30 cycles: 94°C for 20 seconds, 54°C for 30 seconds, 68°C for 20 seconds; final extension at 68°C for 5 minutes. PCR products were purified using Agencort AMPure XP beads (Beckman Coulter) according to the manufacturer’s protocol and purified DNA was eluted in 20-40 μl water.

Ligation-based Illumina library preparation was carried out by the Harvard Biopolymers Facility using an Integex Apollo 324 PrepX ILM kit using Kappa reagents, according to manufacturers’ protocols. For PCR-based Illumina library preparation, samples that had been amplified using one of the barcode-indexed primer sets were subject to a second PCR reaction using an Illumina indexed primer set with regions that were complementary to the barcode lentiviral vector regions (F: JO p50, R: JO p65-p88; Table S5). PCR reaction mixtures contained 100 ng DNA, OneTaq 2x MasterMix (New England Biolabs), and 10 μM each of F/R primers. The following PCR program was used to PCR amplify the template DNA: 94°C for 30 seconds; 15 cycles: 94°C for 20 seconds, 54°C for 30 seconds, 68°C for 20 seconds; final extension at 68°C for 5 minutes. PCR products were purified using Agencort AMPure XP beads (Beckman Coulter) according to the manufacturer’s protocol and purified DNA was eluted in 20-40 μl water.

Barcode specific PCR products with Illumina adaptors were mixed in equimolar ratios and sequenced on an Illumina Miseq at a depth of 20-30×10^6 reads. Reads are first filtered out that do not contain the barcode adaptor sequence ((8 base index)-(17 bases)-ACGCGT-(24 base barcode)-CTGCAG). Samples are divided by 8 base index and a read count table is generated based on all 24-bp barcode sequence present in each read. Each barcode on the read-count table tested for any of the 24-base barcode sequences in the barcode library. A receiver operating characteristic (ROC) curve is generating by diving the 24-bp barcodes into True (present in the barcode library) or False (Absent from the library). Thresholding is performed at the read count that maximized the area under the curve (AUC) from the ROC analysis.

### Detection of barcodes by next-generation sequencing

From genomic DNA preparations, the barcode region of the lentiviral barcode vector was amplified using either primer set JO primer F/R (F: 5’-TGGAGCATGCGCTTTAGCAG; R: ATCGTTTCAGACCCACCTCC-3’) or indexed primer sets (F: JH p05: 5’-ATC GTT TCA GAC CCA CCT CCC-3’; R: JH p11-14 and JH p31-46, Table S2). PCR reaction mixtures contained 250 ng DNA, OneTaq 2x MasterMix (New England Biolabs), and 10 μM each of F/R primers. The following PCR program was used to PCR amplify the template DNA: 94°C for 30 seconds; 30 cycles: 94°C for 20 seconds, 54°C for 30 seconds, 68°C for 20 seconds; final extension at 68°C for 5 minutes. PCR products were purified using Agencort AMPure XP beads (Beckman Coulter) according to the manufacturer’s protocol and purified DNA was eluted in 20-40 μl water.

Ligation-based Illumina library preparation was carried out by the Harvard Biopolymers Facility using an Integex Apollo 324 PrepX ILM kit using Kappa reagents, according to manufacturers’ protocols. Ligation products were PCR amplified using primers against the P5/P7 Illumina adaptor regions. For PCR-based Illumina library preparation, samples that had been amplified using one of the barcode-indexed primer sets were subject to a second PCR reaction using an Illumina indexed primer set with regions that were complementary to the barcode lentiviral vector regions (F: JO p50, R: JO p65-p88, Table S3). PCR reaction mixtures contained 100 ng DNA, OneTaq 2x MasterMix (New England Biolabs), and 10 μM each of F/R primers. The following PCR program was used to PCR amplify the template DNA: 94C for 30 seconds; repeat the following steps for 15 cycles: 94C for 20 seconds, 54C for 30 seconds, 68C for 20 seconds; final extension at 68C for 5 minutes. PCR products were purified using Agencort AMPure XP beads (Beckman Coulter) according to the manufacturer’s protocol and purified DNA was eluted in 20-40 μl water.

Data analysis was performed by identifying index and barcodes in each read by searching for the pattern (8 base index)-(17 bases)-ACGCGT-(24 base barcode)-CTGCAG and counting the index-barcode pairs.

### qPCR-Based Barcode Identification and Analysis

To detect barcodes, gDNA was first pre-amplified by preparing 50μl reactions containing: 500ng gDNA, OneTaq 2x MasterMix (New England Biolabs), and 10μM of each F/R pre-amplification primer to common flanking sequences (Forward: 5’-CGATTAGTGAACGGATCTCG-3’; Reverse: 5’-CCGGTGGATGTGGAATGTG-3’) (Table S4). The following PCR program was used to PCR amplify the template DNA: 94 °C for 30 seconds; 40 cycles at 94 °C for 30 seconds, 52 °C for 30 seconds, 68 °C for 30 seconds; final extension at 68 °C for 5 minutes. The PCR products were purified by Monarch® PCR & DNA Cleanup Kit following the manufacturer’s instructions. Purified PCR products were eluted in 50μl water, and DNA concentration was determined using a Nanodrop 8000 Spectrophotometer (Thermo Fisher). To quantify barcode abundance in any given sample, the purified pre-amplification PCR products were analyzed by qPCR, for which each barcode sequence ^35^ was used as the forward primer and 5’-CCACTTGTGTAGCGCCAAG-3’ was used as the universal reverse primer (Table S4). Each sample was tested for all barcodes by setting up an individual reaction for each barcode primer set per given sample. Each qPCR reaction mixture contained 0.001ng of purified PCR product, 1μM of each F/R primer, and 1X iTaq™ Universal SYBR Green Supermix (Bio-Rad) in 10μL reaction volume. The following program was used: 50 °C for 2 minutes, 94 °C for 10 minutes; 40 cycles: 94 °C for 30 seconds, 60 °C for 30 seconds. Dissociation curves were collected after qPCR, as quality control for qPCR signals to ensure there was a single peak, thus indicating amplification of a single barcode per reaction. The Ct values acquired from the qPCR run were converted to arbitrary units (AU) and the values of the individual barcodes in every sample were summed. The percentage of each barcode present in the sample was calculated by dividing the individual barcode signal by the sum of the total barcode signal.

### Metastasis detection in lung and bone

Total metastasis burden in various tissues was calculated from analysis of gDNA preparations according to the following: The barcode region was pre-amplified from gDNA using forward primer: CGATTAGTGAACGGATCTCG and reverse primer: CCGGTGGATGTGGAATGTG. PCR reaction mixtures contained 100ng DNA, OneTaq 2x MasterMix (New England Biolabs), and 10μM each of F/R primers. The following PCR program was used to amplify the template DNA: 94 °C for 30 seconds; repeat the following steps for 15 cycles: 94 °C for 30 seconds, 52 °C for 30 seconds, 68 °C for 30 seconds; final extension at 68 °C for 5 minutes. The PCR products were purified by Monarch® PCR & DNA Cleanup Kit following manufacturer protocol. Purified PCR products were eluted in 17μl water. Purified PCR products were used in qPCR to quantify total barcode. For each 10μl qPCR reaction, 4μl of the purified PCR products were added to 10μM primers and 2μM probe (forward: TACCGGTTAGTAATGAC; reverse: TAAAGCGCATGCTCCAG; probe: 5’-FAM-AAAAGCGCCTCCCCTACCCGGTAGGTA-3’-Eclipse) and 5μl TaqMan™ Fast Advanced Master Mix (Applied Biosystems™). The following thermocycling program was used: 50 °C for 2 minutes, 95 °C for 2 minutes; repeat the following steps for 40 cycles: 95 °C for 3 seconds, 60 °C for 30 seconds.

### Calibration curve to estimate metastatic burden

We devised a method to extrapolate metastatic burden based on total barcode qPCR signal obtained from different types of tissue. To do so, we first generated calibration curves by spiking known amounts of gDNA isolated from barcoded cells into gDNA isolated from a fixed amount (1 gram) of given tissue from a tumor-free mouse. Based on the estimated amount of gDNA (6 pg) per diploid cell,^56^ we can calculate the estimated barcoded cell number spiked into the given tissue. The serially diluted samples were then subject to PCR-based barcode detection as described above (including pre-amplification, PCR product purification, and qPCR detection using primers and probes, and the TaqMan Fast Advanced Master Mix). Calibration curves were generated by linear regression analysis on log transformed estimated input of barcoded cells and log transformed qPCR signals. Background signal was defined as the qPCR signal of the normal tissue without barcoded tumor cell gDNA. Samples with signal higher than background were defined as positive for metastasis and the metastasis burden interpolated from the calibration curve.

### Statistical Analysis

The data are represented as mean ± SEM, unless otherwise indicated. Data were analyzed by two-way ANOVA with the Sidak multiple comparison correction, one-way ANOVA with Tukey multiple comparison test for significance, or an unpaired two-tailed t test with Welch correction as indicated in the figure legends. All the data were analyzed using GraphPad Prism 7.0. P < 0.05 was considered statistically significant.

## ACKNOWLEDGMENTS

Lentiviral barcode vectors were a generous gift from Dr. Todd R. Golub. We thank Dr. Channing Yu and Lisa Situ for technical assistance. This work was supported by NIH/NHLBI T32 Hematology Training Grant Fellowship to M.S; Landry Cancer Biology Research Fellowship to A.G.M; DOD/CDMRP/BCRP W81XWH-14-1-0191 Era of Hope Scholar Award, NIH/NCI RO1CA166284 Presidential Early Career Award for Scientists and Engineers, and AACR Gertrude B. Elion Cancer Research Award to S.S.M.

## References

1. Wu, A.M., Till, J.E., Siminovitch, L. & McCulloch, E.A. Cytological evidence for a relationship between normal hemotopoietic colony-forming cells and cells of the lymphoid system. J Exp Med 127, 455–464 (1968).

2. Harrison, D.E., Astle, C.M. & Lerner, C. Number and continuous proliferative pattern of transplanted primitive immunohematopoietic stem cells. Proc Natl Acad Sci U S A 85, 822–826 (1988).

3. Capel, B., Hawley, R., Covarrubias, L., Hawley, T. & Mintz, B. Clonal contributions of small numbers of retrovirally marked hematopoietic stem cells engrafted in unirradiated neonatal W/Wv mice. Proc Natl Acad Sci U S A 86, 4564–4568 (1989).

4. Marusyk, A. & Polyak, K. Cancer. Cancer cell phenotypes, in fifty shades of grey. Science 339, 528–529 (2013).

5. Meacham, C.E. & Morrison, S.J. Tumour heterogeneity and cancer cell plasticity. Nature 501, 328–337 (2013).

6. Dagogo-Jack, I. & Shaw, A.T. Tumour heterogeneity and resistance to cancer therapies. Nat Rev Clin Oncol 15, 81–94 (2018).

7. Kreso, A. et al. Variable clonal repopulation dynamics influence chemotherapy response in colorectal cancer. Science 339, 543–548 (2013).

8. Chaffer, C.L. et al. Poised chromatin at the ZEB1 promoter enables breast cancer cell plasticity and enhances tumorigenicity. Cell 154, 61–74 (2013).

9. Castano, Z. et al. IL-1beta inflammatory response driven by primary breast cancer prevents metastasis-initiating cell colonization. Nat Cell Biol 20, 1084–1097 (2018).

10. Campbell, N.R. et al. Cooperation between melanoma cell states promotes metastasis through heterotypic cluster formation. Dev Cell (2021).

11. Navin, N.E. & Hicks, J. Tracing the tumor lineage. Mol Oncol 4, 267–283 (2010).

12. van Galen, P. et al. Single-Cell RNA-Seq Reveals AML Hierarchies Relevant to Disease Progression and Immunity. Cell 176, 1265–1281 e1224 (2019).

13. Acosta, J., Ssozi, D. & van Galen, P. Single-Cell RNA Sequencing to Disentangle the Blood System. Arterioscler Thromb Vasc Biol 41, 1012–1018 (2021).

14. Xu, J. et al. Using single-cell sequencing technology to detect circulating tumor cells in solid tumors. Mol Cancer 20, 104 (2021).

15. Wu, S.Z. et al. A single-cell and spatially resolved atlas of human breast cancers. Nat Genet 53, 1334–1347 (2021).

16. Olive, J.F. et al. Accounting for tumor heterogeneity when using CRISPR-Cas9 for cancer progression and drug sensitivity studies. PLoS One 13, e0198790 (2018).

17. Chaffer, C.L. et al. Normal and neoplastic nonstem cells can spontaneously convert to a stem-like state. Proc Natl Acad Sci U S A 108, 7950–7955 (2011).

18. Janiszewska, M. et al. Subclonal cooperation drives metastasis by modulating local and systemic immune microenvironments. Nat Cell Biol 21, 879–888 (2019).

19. Gerrits, A. et al. Cellular barcoding tool for clonal analysis in the hematopoietic system. Blood 115, 2610–2618 (2010).

20. Lu, R., Neff, N.F., Quake, S.R. & Weissman, I.L. Tracking single hematopoietic stem cells in vivo using high-throughput sequencing in conjunction with viral genetic barcoding. Nat Biotechnol 29, 928–933 (2011).

21. Schepers, K. et al. Dissecting T cell lineage relationships by cellular barcoding. J Exp Med 205, 2309–2318 (2008).

22. Nolan-Stevaux, O. et al. Measurement of Cancer Cell Growth Heterogeneity through Lentiviral Barcoding Identifies Clonal Dominance as a Characteristic of In Vivo Tumor Engraftment. PLoS One 8, e67316 (2013).

23. Nguyen, L.V. et al. DNA barcoding reveals diverse growth kinetics of human breast tumour subclones in serially passaged xenografts. Nat Commun 5, 5871 (2014).

24. Nguyen, L.V. et al. Clonal analysis via barcoding reveals diverse growth and differentiation of transplanted mouse and human mammary stem cells. Cell Stem Cell 14, 253–263 (2014).

25. Porter, S.N., Baker, L.C., Mittelman, D. & Porteus, M.H. Lentiviral and targeted cellular barcoding reveals ongoing clonal dynamics of cell lines in vitro and in vivo. Genome Biol 15, R75 (2014).

26. Bhang, H.E. et al. Studying clonal dynamics in response to cancer therapy using high-complexity barcoding. Nat Med 21, 440–448 (2015).

27. Wagenblast, E. et al. A model of breast cancer heterogeneity reveals vascular mimicry as a driver of metastasis. Nature 520, 358–362 (2015).

28. Yu, C. et al. High-throughput identification of genotype-specific cancer vulnerabilities in mixtures of barcoded tumor cell lines. Nat Biotechnol (2016).

29. Mathis, R.A., Sokol, E.S. & Gupta, P.B. Cancer cells exhibit clonal diversity in phenotypic plasticity. Open Biol 7 (2017).

30. Feldman, D. et al. CloneSifter: enrichment of rare clones from heterogeneous cell populations. BMC Biol 18, 177 (2020).

31. Jin, X. et al. A metastasis map of human cancer cell lines. Nature 588, 331–336 (2020).

32. Oren, Y. et al. Cycling cancer persister cells arise from lineages with distinct programs. Nature (2021).

33. Berthelet, J. et al. The site of breast cancer metastases dictates their clonal composition and reversible transcriptomic profile. Sci Adv 7 (2021).

34. Gutierrez, C. et al. Multifunctional barcoding with ClonMapper enables high-resolution study of clonal dynamics during tumor evolution and treatment. Nature Cancer 2, 758–772 (2021).

35. Peck, D. et al. A method for high-throughput gene expression signature analysis. Genome Biol 7, R61 (2006).

36. Campbell, M.J., Wollish, W.S., Lobo, M. & Esserman, L.J. Epithelial and fibroblast cell lines derived from a spontaneous mammary carcinoma in a MMTV/neu transgenic mouse. In Vitro Cell Dev Biol Anim 38, 326–333 (2002).

37. Miller, F.R. Tumor subpopulation interactions in metastasis. Invasion Metastasis 3, 234–242 (1983).

38. Borowsky, A.D. et al. Syngeneic mouse mammary carcinoma cell lines: two closely related cell lines with divergent metastatic behavior. Clin Exp Metastasis 22, 47–59 (2005).

39. Elenbaas, B. et al. Human breast cancer cells generated by oncogenic transformation of primary mammary epithelial cells. Genes Dev 15, 50–65 (2001).

40. Lehmann, B.D. et al. Identification of human triple-negative breast cancer subtypes and preclinical models for selection of targeted therapies. J Clin Invest 121, 2750–2767 (2011).

41. Nguyen, Q.H. et al. Profiling human breast epithelial cells using single cell RNA sequencing identifies cell diversity. Nat Commun 9, 2028 (2018).

42. Hanahan, D. & Weinberg, R.A. Hallmarks of cancer: the next generation. Cell 144, 646–674 (2011).

43. Wong, K.H., Jin, Y. & Moqtaderi, Z. Multiplex Illumina sequencing using DNA barcoding. Curr Protoc Mol Biol Chapter 7, Unit 7 11 (2013).

44. Mitra, A., Skrzypczak, M., Ginalski, K. & Rowicka, M. Strategies for achieving high sequencing accuracy for low diversity samples and avoiding sample bleeding using illumina platform. PLoS One 10, e0120520 (2015).

45. Aleckovic, M., McAllister, S.S. & Polyak, K. Metastasis as a systemic disease: molecular insights and clinical implications. Biochim Biophys Acta Rev Cancer 1872, 89–102 (2019).

46. Ding, L. et al. Genome remodelling in a basal-like breast cancer metastasis and xenograft. Nature 464, 999–1005 (2010).

47. Gerlinger, M. et al. Intratumor heterogeneity and branched evolution revealed by multiregion sequencing. N Engl J Med 366, 883–892 (2012).

48. Lindstrom, L.S. et al. Clinically used breast cancer markers such as estrogen receptor, progesterone receptor, and human epidermal growth factor receptor 2 are unstable throughout tumor progression. J Clin Oncol 30, 2601–2608 (2012).

49. Luzzi, K.J. et al. Multistep nature of metastatic inefficiency: dormancy of solitary cells after successful extravasation and limited survival of early micrometastases. Am J Pathol 153, 865–873 (1998).

50. Pantel, K., Brakenhoff, R.H. & Brandt, B. Detection, clinical relevance and specific biological properties of disseminating tumour cells. Nat Rev Cancer 8, 329–340 (2008).

51. Hosseini, H. et al. Early dissemination seeds metastasis in breast cancer. Nature (2016).

52. Lucci, A. et al. Circulating tumour cells in non-metastatic breast cancer: a prospective study. Lancet Oncol 13, 688–695 (2012).

53. Bioscience, T. http://www.tmbioscience.com.

54. Luminex http://www.luminexcorp.com.

55. McAllister, S.S. et al. Systemic endocrine instigation of indolent tumor growth requires osteopontin. Cell 133, 994–1005 (2008).

56. Gillooly, J.F., Hein, A. & Damiani, R. Nuclear DNA Content Varies with Cell Size across Human Cell Types. Cold Spring Harb Perspect Biol 7, a019091 (2015).

