## Supplemental Figures and Legends for "SunCatcher: Clonal Barcoding with qPCR-Based Detection Enables Functional Analysis of Live Cells and Generation of Custom Combinations of Cells for Research and Discovery"

#### SUPPLEMENTARY LEGENDS

**Supplementary Figure 1.** Phase contrast microscopic images of indicated HMLER-HR BCs; 20x magnification.

**Supplementary Figure 2. Detection of Luminex DNA barcodes by Sanger sequencing**. **(A)** Primers (JO primer F and JO primer R) were designed upstream and downstream of the barcode region (barcodes were inserted between PstI/MluI restriction sites) to use for PCR amplification prior to Sanger sequencing. **(B)** Agarose gel comparing PCR product yields from PCR reactions using gDNA input extracted using a Qiagen kit (designated as 1; lanes contained reactions with DNA input of 500 ng, 250 ng, and 100 ng, respectively) or the PRISM gDNA extraction method (designated as 2; lanes contained reactions with DNA input of 500 ng, 250 ng, and 100 ng, respectively). Expected amplicon size is ~250 bp (blue arrow). **(C)** Representative chromatogram from Sanger sequencing results of a Luminex barcode amplicon (barcode region is boxed in red.

**Supplementary Figure 3. NGS Detection of BCs.** (**A**) Agarose gel of PCR products following attachment of Illumina adaptors to barcode-index amplicons (Lane 1: 100 bp ladder; Lane 2: no DNA control; Lanes 3-7: PCR products). (**B**) Sequencing read counts for the 3 test samples using the PCR-based Illumina library preparation method. Each library corresponds to a single Illumina adaptor, and the expected barcode pair for each library is indicated.

**Supplementary Figure 4. Characterization of HMLER-HR and 4T1 BC Pools. (A)** Sandplot showing clonal composition of HMLER-HR BC Pool over 6 passages *in vitro*. (**B**) Growth kinetics of HMLER-HR Parental (black) and HMLER-HR BC Pool (grey) over 12 days in culture; n=3 replicates per group; error bars = SD. **(C)** Growth of tumors from 4T1 parental cells (black) and 4T1 BC Pool cells (grey) in Balb/C mice (n=10 per cohort); error bars represent S.D. **(B)** Mass (g) of indicated tumors from (A) at 25-day experimental end point.

**Supplementary Table 1. Collection of Clonally Barcoded Cell Lines**

**Supplementary Table 2. Barcode-index reverse (R) Primer Sequences for NGS-Based Detection.**

**Supplementary Table 3. Illumina Library Preparation Primer Sets.**

**Supplementary Table 4. Oligonucleotide Primer Sequences for qPCR-Based BC Detection.**

**Supplementary Table 5. BC Composition at start and end of injections.** BC composition of the Met1 BC Pool was calculated at the initiation of injections ("pre-injection") and again approximately 2 hr later, when the final injection was performed ("post-injection"). The BC Pool cells were kept in suspension on ice during the injection time period.

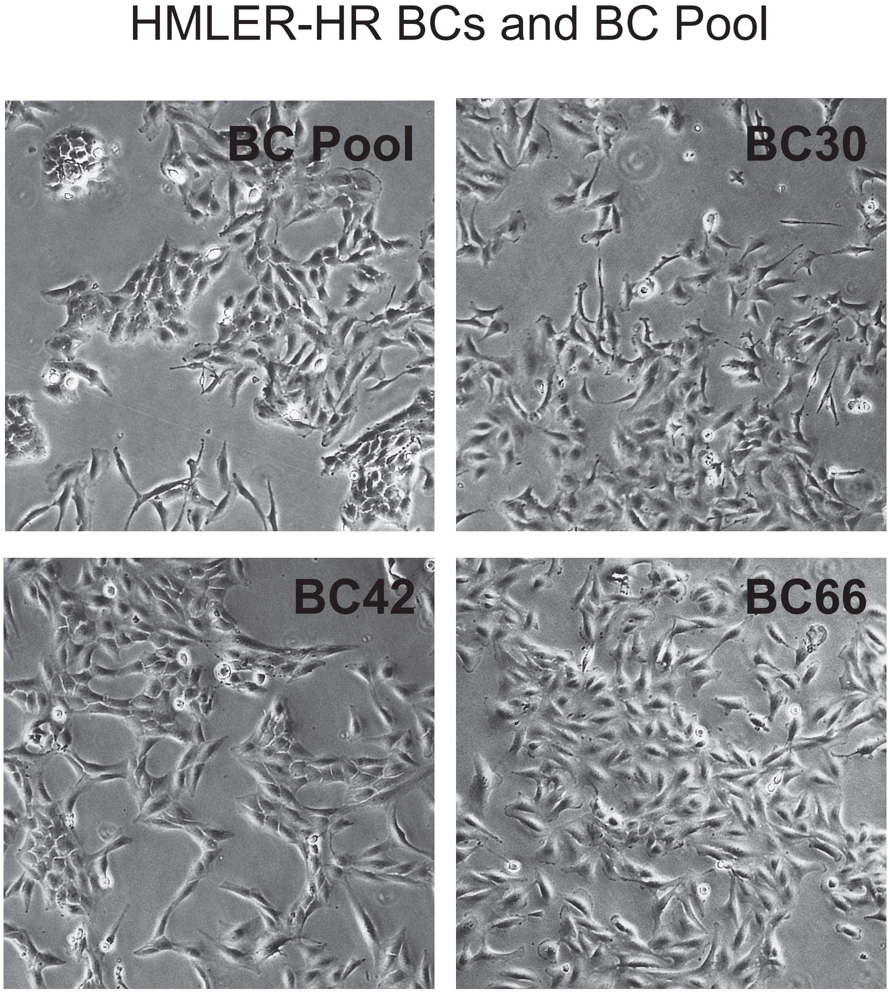


**Supplemental Figure 1. Morphological diversity of HMLER-HR BCs**

### Supplemental Figure 2. Barcode Detection by Sanger Sequencing

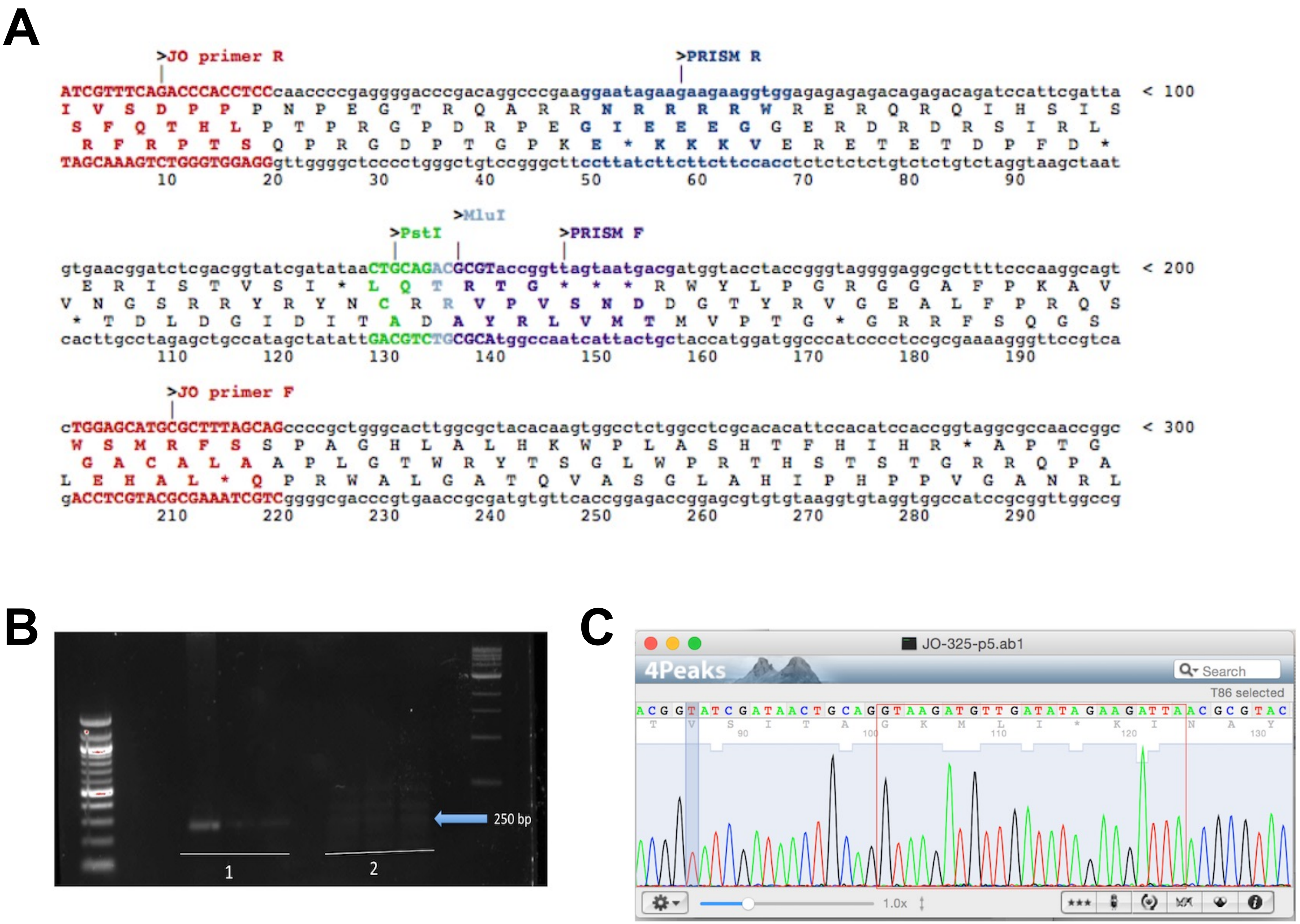

**Supplemental Figure 3. BCs are Accurately Detected by NGS**

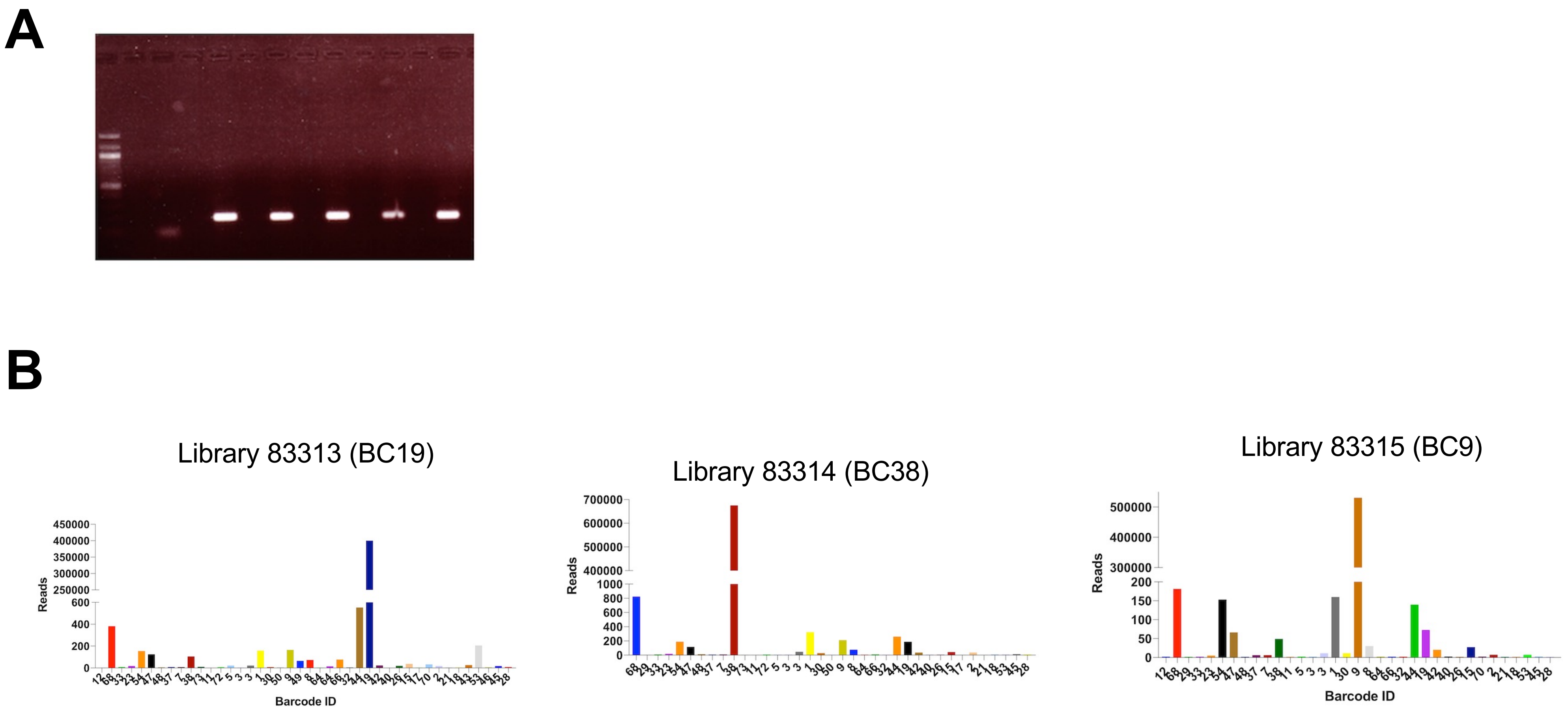

### Supplemental Figure 4. Comparing BC Pools to Parental Cell Lines

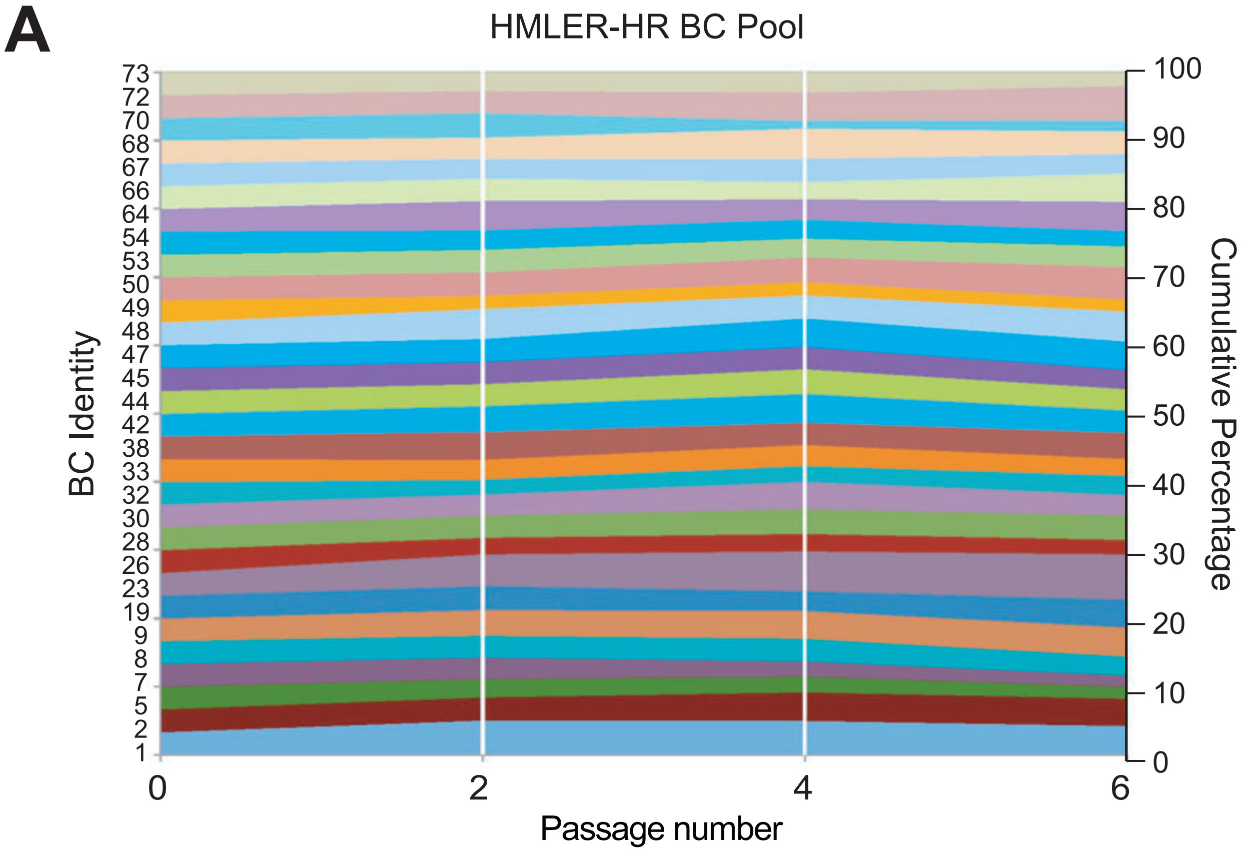


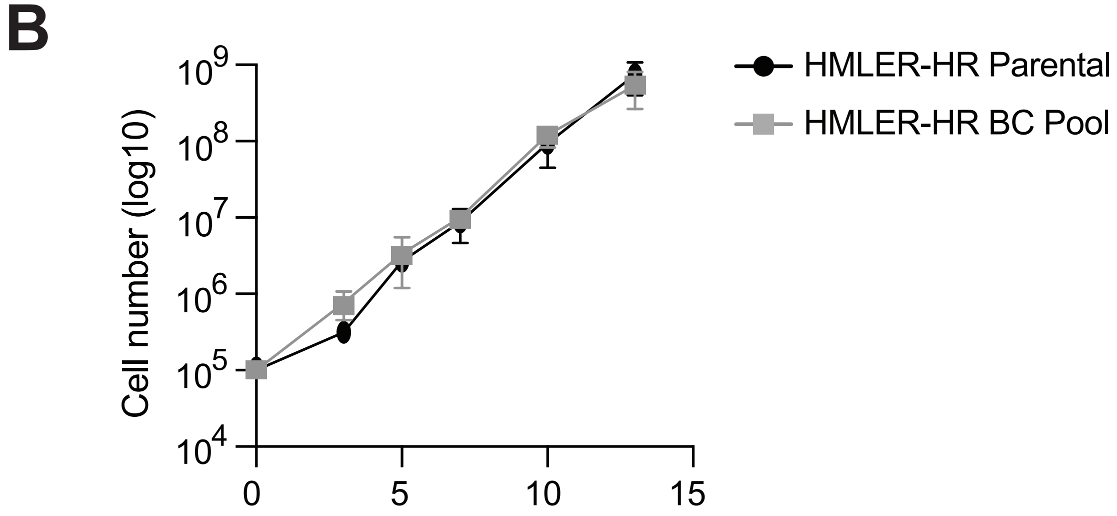


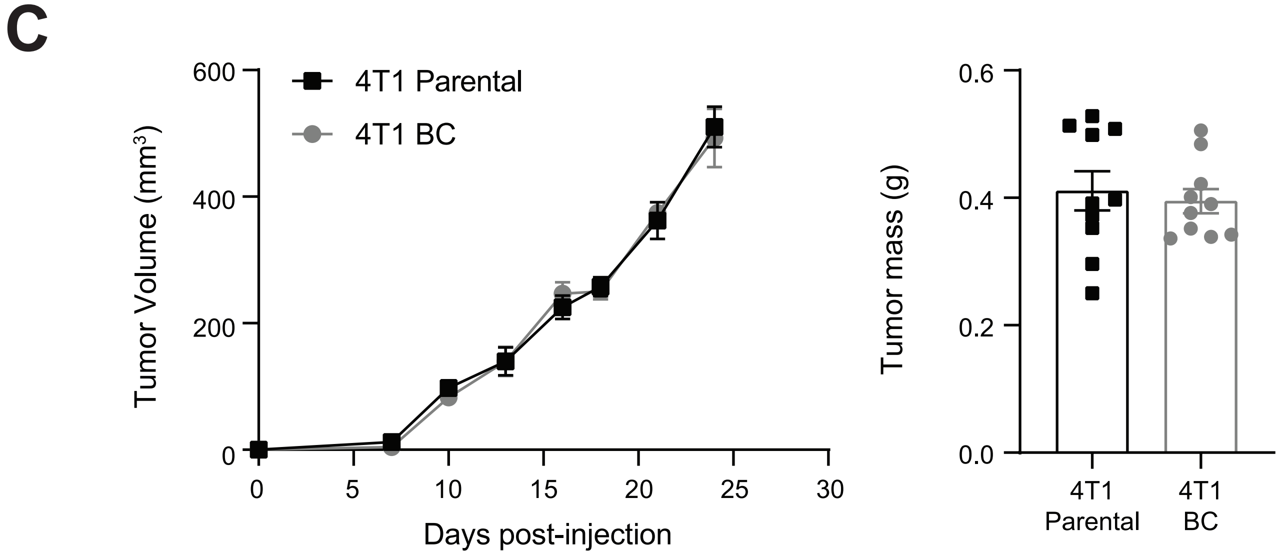

**Supplemental Table 1. Collection of clonally barcoded cell lines**

| BC Pool ID | Species | BC Subtype | No. BCs |
|---|---|---|---|
| Met1 BC Pool | Murine | TNBC | 31 |
| 4T1 BC Pool | Murine | TNBC | 31 |
| MeNeu BC Pool | Murine | Her2+ | 33 |
| HMLER-HR BC Pool | Human | TNBC | 30 |

**Supplemental Table 2. Barcode-index reverse (R) primer sequences**

| Primer | Sequence (Reverse Complement) | Barcode-index # | Barcode Index Sequence |
|---|---|---|---|
| JH p11 | TAGGTGTTCGTCATTACTAACCGG | 1 | AACACCTA |
| JH p12 | AGGCTGGACGTCATTACTAACCGG | 2 | TCCAGCCT |
| JH p13 | GATGCAATCGTCATTACTAACCGG | 3 | ATTGCATC |
| JH p14 | AGCTACGTCGTCATTACTAACCGG | 4 | ACGTAGCT |
| JH p31 | CAGTTAGTCGTCATTACTAACCGG | 5 | ACTAACTG |
| JH p32 | ATCTTAAGCGTCATTACTAACCGG | 6 | CTTAAGAT |
| JH p33 | CTGTTTAACGTCATTACTAACCGG | 7 | TTAAACAG |
| JH p34 | CGAATTCACGTCATTACTAACCGG | 8 | TGAATTCG |
| JH p35 | AATAGGATCGTCATTACTAACCGG | 9 | ATCCTATT |
| JH p36 | TCCTATATCGTCATTACTAACCGG | 10 | ATATAGGA |
| JH p37 | TAGCAACTCGTCATTACTAACCGG | 11 | AGTTGCTA |
| JH p38 | ACTAGCTACGTCATTACTAACCGG | 12 | TAGCTAGT |
| JH p39 | GTTAACTCCGTCATTACTAACCGG | 13 | GAGTTAAC |
| JH p40 | GAATCTAGCGTCATTACTAACCGG | 14 | CTAGATTC |
| JH p41 | GATTCGTACGTCATTACTAACCGG | 15 | TACGAATC |
| JH p42 | GCAATCTTCGTCATTACTAACCGG | 16 | AAGATTGC |
| JH p43 | ACGGTATACGTCATTACTAACCGG | 17 | TATACCGT |
| JH p44 | TGTGACTACGTCATTACTAACCGG | 18 | TAGTCACA |
| JH p45 | TCAGCATTCGTCATTACTAACCGG | 19 | AATGCTGA |
| JH p46 | ATAACGGTCGTCATTACTAACCGG | 20 | ACCGTTAT |

#### Supplemental Table 3. Illumina Library Preparation Primer Sets

| Primer | TruSeq Index | Primer Sequence |
|---|---|---|
| JO p50 | N/A | AATGATACGGCGACCACCGAGATCTACACTCTTTCCCTACACGACGC<br>TCTTCCGATCTGACGGTATCGATATAACTGCAG |
| JO p65 | (1) CGTGAT | CAAGCAGAAGACGGCATACGAGATCGTGATGTGACTGGAGTTCAGA<br>CGTGTGCTCTTCCGATCTCGTCATTACTAACCGG |
| JO p66 | (2) ACATCG | CAAGCAGAAGACGGCATACGAGATACATCGGTGACTGGAGTTCAGA<br>CGTGTGCTCTTCCGATCTCGTCATTACTAACCGG |
| JO p67 | (3) GCCTAA | CAAGCAGAAGACGGCATACGAGATGCCTAAGTGACTGGAGTTCAGA<br>CGTGTGCTCTTCCGATCTCGTCATTACTAACCGG |
| JO p68 | (4) TGGTCA | CAAGCAGAAGACGGCATACGAGATTGGTCAGTGACTGGAGTTCAGA<br>CGTGTGCTCTTCCGATCTCGTCATTACTAACCGG |
| JO p69 | (5) CACTGT | CAAGCAGAAGACGGCATACGAGATCACTGTGTGACTGGAGTTCAGAC<br>GTGTGCTCTTCCGATCTCGTCATTACTAACCGG |
| JO p70 | (6) ATTGGC | CAAGCAGAAGACGGCATACGAGATATTGGCGTGACTGGAGTTCAGA<br>CGTGTGCTCTTCCGATCTCGTCATTACTAACCGG |
| JO p71 | (7) GATCTG | CAAGCAGAAGACGGCATACGAGATGATCTGGTGACTGGAGTTCAGA<br>CGTGTGCTCTTCCGATCTCGTCATTACTAACCGG |
| JO p72 | (8) TCAAGT | CAAGCAGAAGACGGCATACGAGATTCAAGTGTGACTGGAGTTCAGAC<br>GTGTGCTCTTCCGATCTCGTCATTACTAACCGG |
| JO p73 | (9) CTGATC | CAAGCAGAAGACGGCATACGAGATCTGATCGTGACTGGAGTTCAGAC<br>GTGTGCTCTTCCGATCTCGTCATTACTAACCGG |
| JO p74 | (10) AAGCTA | CAAGCAGAAGACGGCATACGAGATAAGCTAGTGACTGGAGTTCAGAC<br>GTGTGCTCTTCCGATCTCGTCATTACTAACCGG |
| JO p75 | (11) GTAGCC | CAAGCAGAAGACGGCATACGAGATGTAGCCGTGACTGGAGTTCAGA<br>CGTGTGCTCTTCCGATCTCGTCATTACTAACCGG |
| JO p76 | (12) TACAAG | CAAGCAGAAGACGGCATACGAGATTACAAGGTGACTGGAGTTCAGAC<br>GTGTGCTCTTCCGATCTCGTCATTACTAACCGG |
| JO p77 | (13) TTGACT | CAAGCAGAAGACGGCATACGAGATTTGACTGTGACTGGAGTTCAGAC<br>GTGTGCTCTTCCGATCTCGTCATTACTAACCGG |
| JO p78 | (14) GGAACT | CAAGCAGAAGACGGCATACGAGATGGAACTGTGACTGGAGTTCAGA<br>CGTGTGCTCTTCCGATCTCGTCATTACTAACCGG |
| JO p79 | (15) TGACAT | CAAGCAGAAGACGGCATACGAGATTGACATGTGACTGGAGTTCAGAC<br>GTGTGCTCTTCCGATCTCGTCATTACTAACCGG |
| JO p80 | (16) GGACGG | CAAGCAGAAGACGGCATACGAGATGGACGGGTGACTGGAGTTCAGA<br>CGTGTGCTCTTCCGATCTCGTCATTACTAACCGG |
| JO p81 | (18) GCGGAC | CAAGCAGAAGACGGCATACGAGATGCGGACGTGACTGGAGTTCAGA<br>CGTGTGCTCTTCCGATCTCGTCATTACTAACCGG |
| JO p82 | (19) TTTCAC | CAAGCAGAAGACGGCATACGAGATTTTCACGTGACTGGAGTTCAGAC<br>GTGTGCTCTTCCGATCTCGTCATTACTAACCGG |
| JO p83 | (20) GGCCAC | CAAGCAGAAGACGGCATACGAGATGGCCACGTGACTGGAGTTCAGA<br>CGTGTGCTCTTCCGATCTCGTCATTACTAACCGG |
| JO p84 | (21) CGAAAC | CAAGCAGAAGACGGCATACGAGATCGAAACGTGACTGGAGTTCAGA<br>CGTGTGCTCTTCCGATCTCGTCATTACTAACCGG |
| JO p85 | (22) CGTACG | CAAGCAGAAGACGGCATACGAGATCGTACGGTGACTGGAGTTCAGA<br>CGTGTGCTCTTCCGATCTCGTCATTACTAACCGG |
| JO p86 | (23) CCACTC | CAAGCAGAAGACGGCATACGAGATCCACTCGTGACTGGAGTTCAGA<br>CGTGTGCTCTTCCGATCTCGTCATTACTAACCGG |
| JO p87 | (25) ATCAGT | CAAGCAGAAGACGGCATACGAGATATCAGTGTGACTGGAGTTCAGAC<br>GTGTGCTCTTCCGATCTCGTCATTACTAACCGG |
| JO p88 | (27) AGGAAT | CAAGCAGAAGACGGCATACGAGATAGGAATGTGACTGGAGTTCAGA<br>CGTGTGCTCTTCCGATCTCGTCATTACTAACCGG |

#### Supplemental Table 4. qPCR primer sequence for BC detection

| Oligo ID | Sequence(5′-3′) | Length(bp) | GC content | Tm(°C) |
|---|---|---|---|---|
| BC1 | GATTTGTATTGATTGAGATTAAAG | 24 | 0.25 | 50.1 |
| BC2 | TGATTGTAGTATGTATTGATAAAG | 24 | 0.25 | 50.1 |
| BC3 | GATTGTAAGATTTGATAAAGTGTA | 24 | 0.25 | 50.1 |
| BC4 | GATTTGAAGATTATTGGTAATGTA | 24 | 0.25 | 50.1 |
| BC5 | GATTGATTATTGTGATTTGAATTG | 24 | 0.25 | 50.1 |
| BC7 | ATTGGTAAATTGGTAAATGAATTG | 24 | 0.25 | 50.1 |
| BC8 | GTAAGTAATGAATGTAAAAGGATT | 24 | 0.25 | 50.1 |
| BC9 | GTAAGATGTTGATATAGAAGATTA | 24 | 0.25 | 50.1 |
| BC11 | GATTAAAGTGATTGATGATTTGTA | 24 | 0.25 | 50.1 |
| BC13 | TTAGTGAAGAAGTATAGTTTATTG | 24 | 0.25 | 50.1 |
| BC14 | AAAGTATAGTAAGATGTATAGTAG | 24 | 0.25 | 50.1 |
| BC15 | TGAATTGATGAATGAATGAAGTAT | 24 | 0.25 | 50.1 |
| BC16 | TGATGATTTGAATGAAGATTGATT | 24 | 0.25 | 50.1 |
| BC17 | TGATAAAGTGATAAAGGATTAAAG | 24 | 0.25 | 50.1 |
| BC18 | TGATTTGAGTATTTGAGATTTTGA | 24 | 0.25 | 50.1 |
| BC19 | GTATTTGAGTAAGTAATTGATTGA | 24 | 0.25 | 50.1 |
| BC20 | GATTGTATTGAAGTATTGTAAAAG | 24 | 0.25 | 50.1 |
| BC21 | TGATTTGAGATTAAAGAAAGGATT | 24 | 0.25 | 50.1 |
| BC22 | TGATTGAATTGAGTAAAAAGGATT | 24 | 0.25 | 50.1 |
| BC23 | AAAGTTGAGATTTGAATGATTGAA | 24 | 0.25 | 50.1 |
| BC24 | GTATTGTATTGAAAAGGTAATTGA | 24 | 0.25 | 50.1 |
| BC25 | TGAAGATTTGAAGTAATTGAAAAG | 24 | 0.25 | 50.1 |
| BC26 | TGAAAAAGTGTAGATTTTGAGTAA | 24 | 0.25 | 50.1 |
| BC27 | AAAGTTGAGTATTGATTTGAAAAG | 24 | 0.25 | 50.1 |
| BC28 | TTGATAATGTTTGTTTGTTTGTAG | 24 | 0.25 | 50.1 |
| BC29 | AAAGAAAGGATTTGTAGTAAGATT | 24 | 0.25 | 50.1 |
| BC30 | GTAAAAAGAAAGGTATAAAGGTAA | 24 | 0.25 | 50.1 |
| BC31 | GATTAAAGTTGATTGAAAAGTGAA | 24 | 0.25 | 50.1 |
| BC32 | GTAGATTAGTTTGAAGTGAATAAT | 24 | 0.25 | 50.1 |
| BC33 | AAAGGATTAAAGTGAAGTAATTGA | 24 | 0.25 | 50.1 |
| BC36 | ATTGATTGTGAATGAAATGAATTG | 24 | 0.25 | 50.1 |
| BC37 | ATTGAAAGATGAAAAGATGAAAAG | 24 | 0.25 | 50.1 |
| BC38 | ATTGTTGAAAAGTGTAATGATTGA | 24 | 0.25 | 50.1 |
| BC40 | TAATGTTGTGAATAATGTAGAAAG | 24 | 0.25 | 50.1 |
| BC42 | GTTTATAGTGAAATATGAAGATAG | 24 | 0.25 | 50.1 |
| BC43 | TGTAATGAGTATTGTAATTGAAAG | 24 | 0.25 | 50.1 |
| BC44 | GTATAAAGAAAGATTGGTAAATGA | 24 | 0.25 | 50.1 |
| BC45 | TTGAGTAATTGAATTGTGAAATGA | 24 | 0.25 | 50.1 |
| BC46 | TGTATTGAATGAATTGTTGATGTA | 24 | 0.25 | 50.1 |
| BC47 | ATTATGAAGTAAGTTAATGAGAAG | 24 | 0.25 | 50.1 |
| BC48 | ATTATTGAGATGTGAAGTTTGTTT | 24 | 0.25 | 50.1 |
| BC50 | GTAAATGATGATATTGGTATATTG | 24 | 0.25 | 50.1 |
| BC52 | ATTGTGAAGTATAAAGATGATTGA | 24 | 0.25 | 50.1 |
| BC53 | TGTAGAAGATGAGATGTATAATTA | 24 | 0.25 | 50.1 |
| BC54 | ATGAATTGAAAGTGATTGAAAAAG | 24 | 0.25 | 50.1 |
| BC57 | GTAATGATAAAGATGATGATATTG | 24 | 0.25 | 50.1 |
| BC64 | GTAAGTAGTAATTTGAATATGTAG | 24 | 0.25 | 50.1 |
| BC65 | AAAGGTAAGATTATTGATGAAAAG | 24 | 0.25 | 50.1 |
| BC66 | GTAGATAGTATAGTTGTAATGTTA | 24 | 0.25 | 50.1 |
| BC67 | GATTTGTAATTGTTGAGTAAATGA | 24 | 0.25 | 50.1 |
| BC68 | AAAGAAAGATTGTTGAGATTATGA | 24 | 0.25 | 50.1 |
| BC69 | GATGTGAATGTAATATGTTTATAG | 24 | 0.25 | 50.1 |
| BC70 | TGATATGAATTGGATTATTGGTAT | 24 | 0.25 | 50.1 |
| BC71 | ATGAATTGATTGGATTGTAATGAT | 24 | 0.25 | 50.1 |
| BC72 | GATTATTGGATTAAAGGTAAATGA | 24 | 0.25 | 50.1 |
| BC73 | ATTGTTGAATTGATGAGATTTGAT | 24 | 0.25 | 50.1 |
| BC74 | TGAAATTAGTTTGTAAGATGTGTA | 24 | 0.25 | 50.1 |
| PreAmp_F | CGATTAGTGAACGGATCTCG | 20 | 0.50 | 60.0 |
| PreAmp_R | CCGGTGGATGTGGAATGTG | 19 | 0.58 | 60.0 |
| BCUniversal | CCACTTGTGTAGCGCCAAG | 19 | 0.58 | 60.0 |

**Supplemental Table 5. Contribution (%) of Each BC to BC Pool at Time of Injection and After the Last Mouse was Injected**

| | % Composition | |
|---|---|---|
| BC | Pre-Injection | Post-Injection |
| 2 | 0.7459 | 0.8200 |
| 3 | 0.2218 | 0.1703 |
| 5 | 0.1089 | 0.1180 |
| 7 | 4.3510 | 4.7488 |
| 8 | 2.0571 | 2.6568 |
| 11 | 1.5696 | 1.5971 |
| 14 | 0.2405 | 0.2442 |
| 17 | 0.0207 | 0.0203 |
| 18 | 1.0353 | 1.1722 |
| 19 | 2.8757 | 2.9717 |
| 20 | 8.3150 | 9.6790 |
| 21 | 0.4561 | 0.5436 |
| 22 | 1.2243 | 1.5013 |
| 23 | 1.7757 | 1.8088 |
| 25 | 0.7883 | 0.5301 |
| 26 | 0.6720 | 0.6867 |
| 28 | 0.3914 | 0.2831 |
| 30 | 1.0937 | 1.1002 |
| 31 | 1.0464 | 0.9739 |
| 33 | 0.3400 | 0.3054 |
| 36 | 2.7291 | 2.2966 |
| 37 | 0.0587 | 0.0698 |
| 40 | 0.2769 | 0.2854 |
| 42 | 1.3537 | 1.3695 |
| 43 | 4.1184 | 5.6051 |
| 44 | 0.8892 | 0.8828 |
| 45 | 13.1404 | 11.0982 |
| 48 | 0.1662 | 0.1425 |
| 53 | 25.6185 | 24.6964 |
| 67 | 20.5118 | 19.9264 |
| 73 | 1.8079 | 1.6960 |
